# Cell-Sized Lipid Vesicles as Artificial Antigen-Presenting Cells for Antigen-Specific T Cell Activation

**DOI:** 10.1101/2022.12.02.518936

**Authors:** Jui-Yi Chen, Sudhanshu Agrawal, Hsiu-Ping Yi, Derek Vallejo, Anshu Agrawal, Abraham Lee

**Affiliations:** Biomedical Engineering, University of California, Irvine, CA 92617, USA; School of Medicine, University of California, Irvine, CA 92617, USA

**Keywords:** droplet generation, microfluidic, artificial antigen-presenting cell, T cell activation, immunotherapy

## Abstract

In this study, efficient T cell activation is demonstrated using cell-sized artificial antigen-presenting cells (aAPCs) with protein-conjugated bilayer lipid membranes that mimic biological cell membranes. The highly uniform aAPCs are generated by a facile method based on standard droplet microfluidic devices. These aAPCs are able to activate the T cells in peripheral blood mononuclear cells (PBMCs), showing a 28-fold increase in IFNγ secretion, a 233-fold increase in antigen-specific CD8 T cells expansion, and a 16-fold increase of CD4 T cell expansion. The aAPCs do not require repetitive boosting or additional stimulants and can function at a relatively low aAPC-to-T cell ratio (1-to-17). The research presents strong evidence that the surface fluidity and size of the aAPCs are critical to the effective formation of immune synapses essential for T cell activation. The findings demonstrate that the microfluidic-generated aAPCs can be instrumental in investigating the physiological conditions and mechanisms for T cell activation. Finally, this method demonstrates the feasibility of customizable aAPCs for a cost-effective off-the-shelf approach to immunotherapy.

## 1. Introduction

Immunotherapy utilizes the activation or suppression of immune responses to treat diseases such as autoinflammation, autoimmune diseases, and cancer.^[1]^ One type of immunotherapy that is widely applied to treat cancer is adoptive cell transfer (ACT), which extracts the immune cells from patients, engineers the cells to enhance immunogenicity, expands cells in large numbers, and then reinfuses them to the patients. The earliest T cell-based ACT was demonstrated by tumor-infiltrating lymphocyte (TIL) therapy, which utilized autologous T cells to treat patients with metastatic melanoma.^[2]^ Following that, genetic engineering further boosted the development of T cell receptor (TCR) therapy and chimeric antigen receptor (CAR) T cell therapy.^[3]^ While these treatments have shown remarkable results, their popularity was limited by the insufficient expansion of potent T cells.^[4]^ It typically requires weeks to months to expand the T cells with antigen-specificity,^[5]^ and some patients cannot survive long enough to wait for the processing time. Therefore, there is a great demand for an efficient and robust technology to produce high-quality T cells.

In the human body, T cells are activated by antigen-presenting cells (APCs), especially dendritic cells (DCs). While DCs were first explored to expand T cells, the use of DCs is still challenging due to several reasons. Firstly, DCs exhibit suppressive phenotypes in cancer that will downregulate T cell activation ^[6, 7]^. Secondly, their culturing process is labor-intensive and requires a combination of expensive cytokines.^[8, 9]^ Thirdly, the low population of DCs makes it difficult for downstream processing. Finally, the individual variation in DC quantity and quality renders inconsistent results of T cell expansion.^[4]^ Cell lines, such as K562 and NIH/3T3, have been employed.^[10–12]^ However, they have not been widely accepted due to the concern of infusing malignant clones into cancer patients.^[4]^ These limitations have led to a great need for substituting natural APCs with artificial APCs (aAPCs) (**Figure 1**).

**Figure 1.**
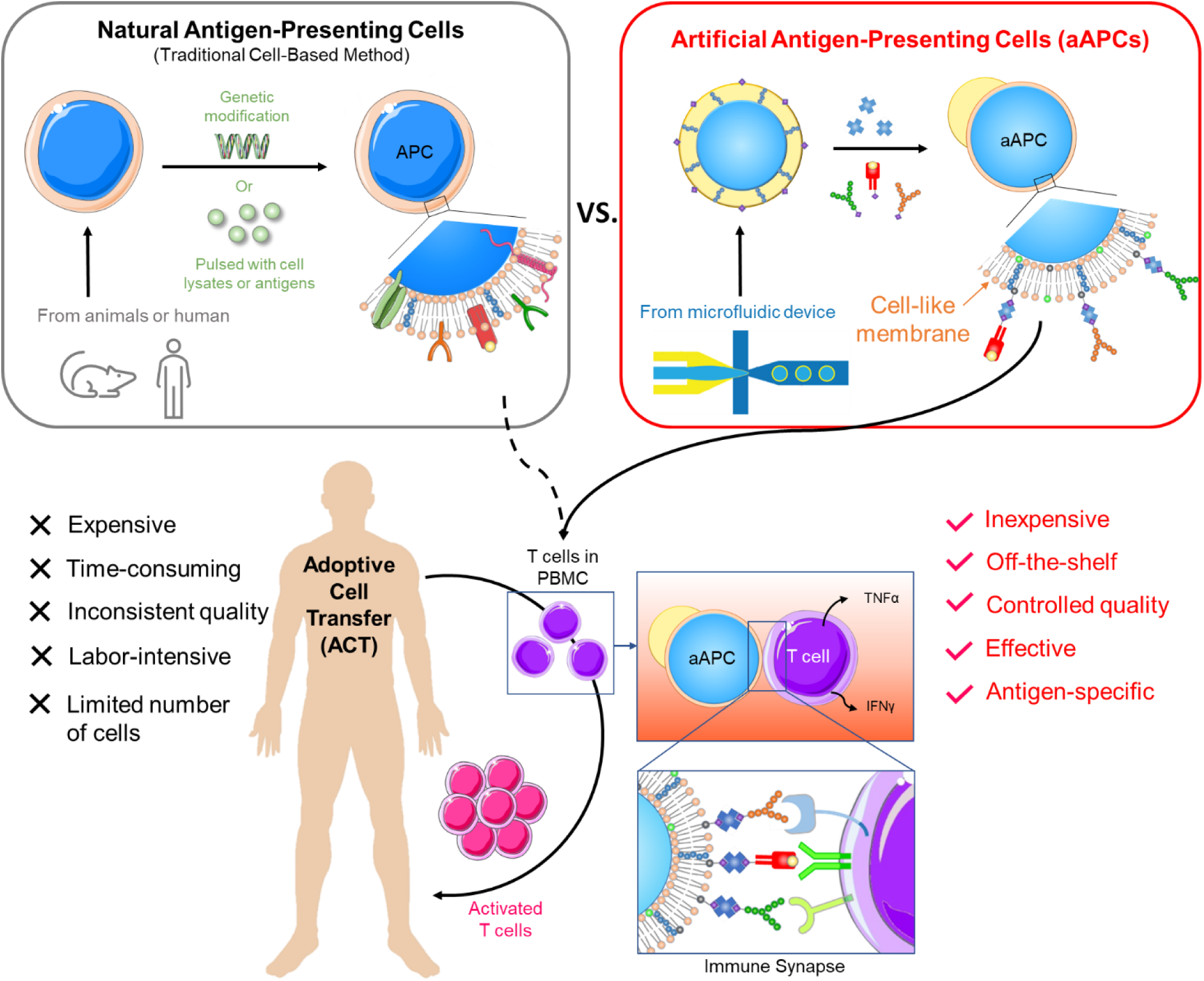
The schematic illustration of adoptive cell transfer (ACT) using natural APCs and artificial APCs (aAPCs). Natural APCs are prepared by 1) deriving dendritic cells (DCs) from monocytes and pulsing them with cell lysates or antigens or 2) genetically modifying the cell lines, such as K562 myelogenous leukemia cells or NIH/3T3 fibroblast cells, to give them antigen-presentation functions. The process is long, and the quality is inconsistent due to the variation in cells. aAPCs proposed in this research are made by dewetting excessive oil of a double-emulsion droplet (DED), followed by conjugating the surface proteins. The aAPCs can be produced in large quantities in a short time at a low cost. The as-prepared aAPC has a fluidic lipid bilayer that makes it easy to form an immune synapse with T cells, resulting in effective T cell activation.

Artificial APCs (aAPCs) activate T cells through the engagement of multiple ligands and receptors on each cell. Once the T cell receptor (TCR) on a T cell recognizes the peptide-loaded MHC (pMHC) on the APC, the bull’s-eye structure starts to form at the cellular interface, with TCRs and CD28 accumulating at the center (named as central supramolecular activation cluster, cSMAC) and LFA-1 (lymphocyte function-associated antigen 1) surrounding the central clusters (named as peripheral SMAC or pSMAC).^[13]^ Since the receptors on the T cell are highly structured, aAPCs with high membrane fluidity are needed to maximize the bindings of TCR, CD28, and LFA-1, forming a sturdy immune synapse.^[14]^ Membrane fluidity and size are critical as the TCRs can be triggered more effectively when the aAPCs are in the size range of a cell (10-25 µm). ^[15]^ This can be explained by the mechano-sensing nature of TCRs. After the TCR-pMHC bonding has formed, the mechanical forces (compression, tension, and shear stress) generated by the relative movement between T cells and dendritic cells can induce the TCR transduction pathway. ^[16, 17]^ Therefore, the adequate size of an aAPC ensures the sufficient generation of mechanical forces, thus enhancing the activation of T cells.

Various materials have been used to construct aAPCs. For instance, magnetic beads (4.5 µm) coated with human leukocyte antigen–immunoglobulin fusion protein (HLA-Ig) and anti-CD28 antibodies have been applied to activate antigen-specific T cells. ^[18]^ Polymeric microspheres, such as polystyrene and poly lactic-co-glycolic acid (PLGA) particles, ^[19, 20]^ have also been extensively used. While the polymers are softer and more fluid, the entanglement of the long polymeric chain and the relatively high melting point renders a rigid surface and thus restricting the movement of ligands.^[14]^ By far, a fluid surface to mimic cell membranes can be best achieved using lipid membranes or lipid-containing solutions.

Liposomes have been of great interest for aAPC production as they can recapitulate the cell membrane fluidity. ^[21, 22]^ Yet, conventional methods for liposome production, such as thin-film hydration, ^[23]^ electroformation, ^[24]^ ethanol injection, ^[25]^ and reverse-phase evaporation ^[26]^ are unable to produce monodisperse, cell-sized (10-25 µm in diameter) liposomes. Extrusion method can reduce multi-layer liposomes to unilamellar (single-layer) liposomes with high monodispersity.^[27]^ However, this method typically produces liposomes less than 200 nm in diameter, about two orders of magnitude lower than the desired range. Droplet microfluidics offers an alternative way to generate cell-sized liposomes with high uniformity. ^[28–32]^ However, these methods are mostly restricted by low yield and short storage time.

This study presents a facile method to produce monodispersed, cell-sized aAPCs with a fluid membrane conjugated with immune synapse ligands. Double emulsion droplets (DEDs) are generated using a flow-focusing microfluidic device to form droplets with an aqueous inner core and a lipid-carrying oil shell. DEDs are then converted into single-compartment multisomes (SCMs) by a dewetting process that transforms the oil shell into a bilayer lipid membrane and an oil cap. ^[33, 34]^ SCMs are then conjugated with immune-synapse antibodies, namely anti-CD28, anti-LFA-1, and pMHC, forming aAPCs with a bilayer lipid membrane termed aAPC-BLMs. On the other hand, DEDs that are directly conjugated with the same antibodies while still with an oil shell are termed aAPC-shells. aAPC-shells are constructed along with aAPC-BLMs to study whether the fluid lipid membrane that mimics a real cell membrane can improve T cell activation. Since cytokine secretion is upregulated when naïve T cells are activated by APCs, the cytokine levels are measured to quantify T cell activation by aAPCs as well. Specifically, TNFα, IFNγ, IL-2, and IL-10 are the cytokines selected for our experiments. TNFα is measured since it is secreted and received by T cells to promote activation and proliferation. ^[35]^ On the other hand, secretion of IFNγ by effector CD4 and CD8 T cells are known to improve the recognition of tumor cells. ^[36]^ In general, the enhanced secretion of TNFα and IFNγ are positively correlated with the portion of CD4 Th1 cells and CD8 Tc1 cells both of which favor the killing of tumor. ^[36, 37]^ IL-2 is highly secreted in activated T cells to promote proliferation. Finally, IL-10 downregulates the anti-tumor response by restricting T cell activation. Therefore, the activation of T cells is analyzed by measuring these cytokines (TNFα, IFNγ, IL-2, and IL-10). NYESO-1 peptide is chosen to demonstrate the potency of aAPCs since it is a unique class of tumor-associated antigen that has restricted expression in normal somatic tissues and is overly expressed in many types of tumors. ^[38, 39]^ Furthermore, it has also been a promising antigen for vaccine studies and adoptive immunotherapy. ^[40, 41]^

Various control groups are designed in this study to investigate how aAPC sizes and their surface fluidity contribute to efficient T cell activation. The control groups selected for the study include oil drops, polystyrene beads, and free molecules. This is the first demonstration of effective antigen-specific T cell activation by aAPCs based on highly-uniform, cell-sized lipid vesicles. In summary, we present a flexible and versatile process to optimize artificial antigen-presenting cells for physiological interaction with biological cells that shows promise for applications in customizable personal medicine treatments.

## 2. Results and Discussions

### 2.1. Generation of DEDs

Double emulsion droplets (DEDs) were generated using a single-step flow-focusing device (Figure 2a-c, f&g) fabricated by photolithography (S 1). The device allows for stable droplet generation to go on for more than 3 hours, with a droplet generation rate of 250 drops s-1. The DEDs have an average size of 22.36 ± 1.09 µm (CV = 4.90 %, PDI = 0.0024, N = 302, Figure 2i), which is highly monodisperse (PDI < 0.05) and is close to the nominal size of a cell. ^[42, 43]^ In addition, DEDs have high stability, with 90 % remaining at room temperature after one month (S 5). The high stability allows DEDs to be stored for a long time and then be converted into SCMs upon usage.^[34]^ A detailed description of the channel improvement can be found in the supporting information (S 1-S 5).

**Figure 2.**
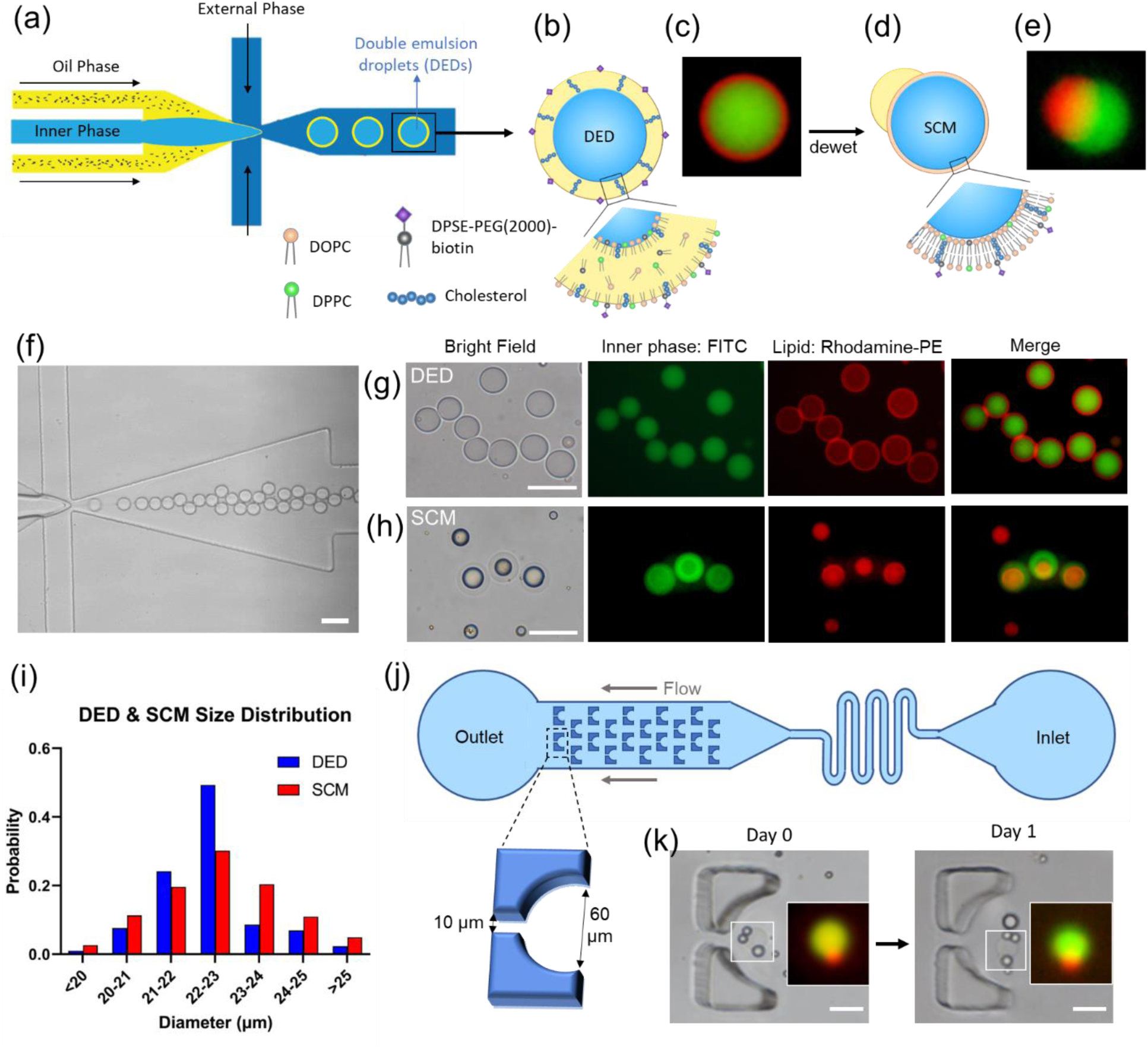
Generation of double emulsion droplets (DEDs) and single-compartment multisomes (SCMs). (a) Illustration of DED generation on a flow-focusing microfluidic device. (b) Illustration figure and (c) fluorescence image of DED. (c) Images of DEDs generated with 0.05 mol% rhodamine-PE lipids (red) encapsulating FITC (green). (d) Illustration figure and (e) fluorescence image of SCMs converted from DEDs, showing SCMs have an oil cap and a lipid bilayer membrane. (f) Generation of DEDs. (g) Images of DEDs. (h) Images of SCMs. (i) Size distribution of DED and SCM. (j) Illustration of the trapping array. (k) SCMs are stable in the trapping array filled with RPMI + 10% FBS after one day. Scale bar = 50 µm.

### 2.2. Convert DEDs into SCMs, which stay stable in a RPMI-filled trapping array overnight

To generate a cell-like lipid bilayer membrane, the DEDs were converted into SCMs by the dewetting mechanism. As presented by Vallejo et al.,^[34]^ the DEDs can be kept for months and then be transformed into SCMs by controlling the interfacial energies. DEDs have a thin oil shell, which looks like a red ring under a fluorescent microscope (Figure 2g). SCMs were generated by immersing DEDs in 1X PBS (Figure 2d, e, and h). The excessive oil was dewetted, forming a lipid bilayer with an oil cap in 10 minutes. The lipid bilayer in SCMs is red (rhodamine-PE) but is too thin to be visualized clearly. The oil cap can be seen on the side of a SCM (Figure 2e, S 10b) or on top of it (Figure 2h). SCMs have an average size of 22.54 ± 1.39 µm (CV = 6.20%, PDI = 0.0038, N = 265, Figure 2i), similar in size to DEDs. SCMs are slightly more polydisperse possibly due to different degrees of dewetting. The SCMs are, however, highly monodisperse compared to the liposomes generated by conventional methods (e.g., thin-film hydration and electroformation),^[24]^ ^[44, 45]^ where PDI falls between 0.1-0.2.

A trapping array was designed to observe the stability of SCMs (Figure 2j). DEDs were loaded into the reservoir inlet, and a syringe was pulled from the outlet. The trap has a circular opening of 60 µm in diameter and a 10 µm gap, allowing the flow to pass through and immobilize the droplet. DEDs were captured and transformed into SCMs on-chip, and then RPMI with 10 % FBS was flown into the channel. SCMs were still stable after the RPMI flowed in, and their structure remained intact after one day (Figure 2k).

### 2.3. Verification of unilamellarity

This section aims to prove that the dewetted DEDs have a lipid bilayer (or unilamellar) membrane. The unilamellarity is proven by three methods: osmotic shock, membrane protein insertion, and identifying the inner and outer leaflets.

#### 2.3.1. Osmotic shock

A lipid bilayer is a semi-permeable membrane that blocks the ions and large molecules but allows the small molecules and water to pass through. When placed in a hypertonic solution, the SCMs shrink as the water exits the droplet. The permeability of the membrane can be calculated by measuring the radius of SCM over a period of time, according to equation (b). ^[46, 47]^ In our experiment, the membrane permeability to water was calculated to be 53.6 ± 3.4 μm s-1 (Figure 3), which is well within the range of 25-150 μm s-1 for DOPC bilayers. ^[48, 49]^ A similar test conducted on DEDs resulted in values between 2-14 μm s-1 (data not shown), confirming that any water passing through the oil cap can be considered minimal, compared to the rate of water passing through the membrane. Therefore, the measurement accurately reflects that the SCMs have the permeability of a lipid bilayer vesicle.

**Figure 3.**
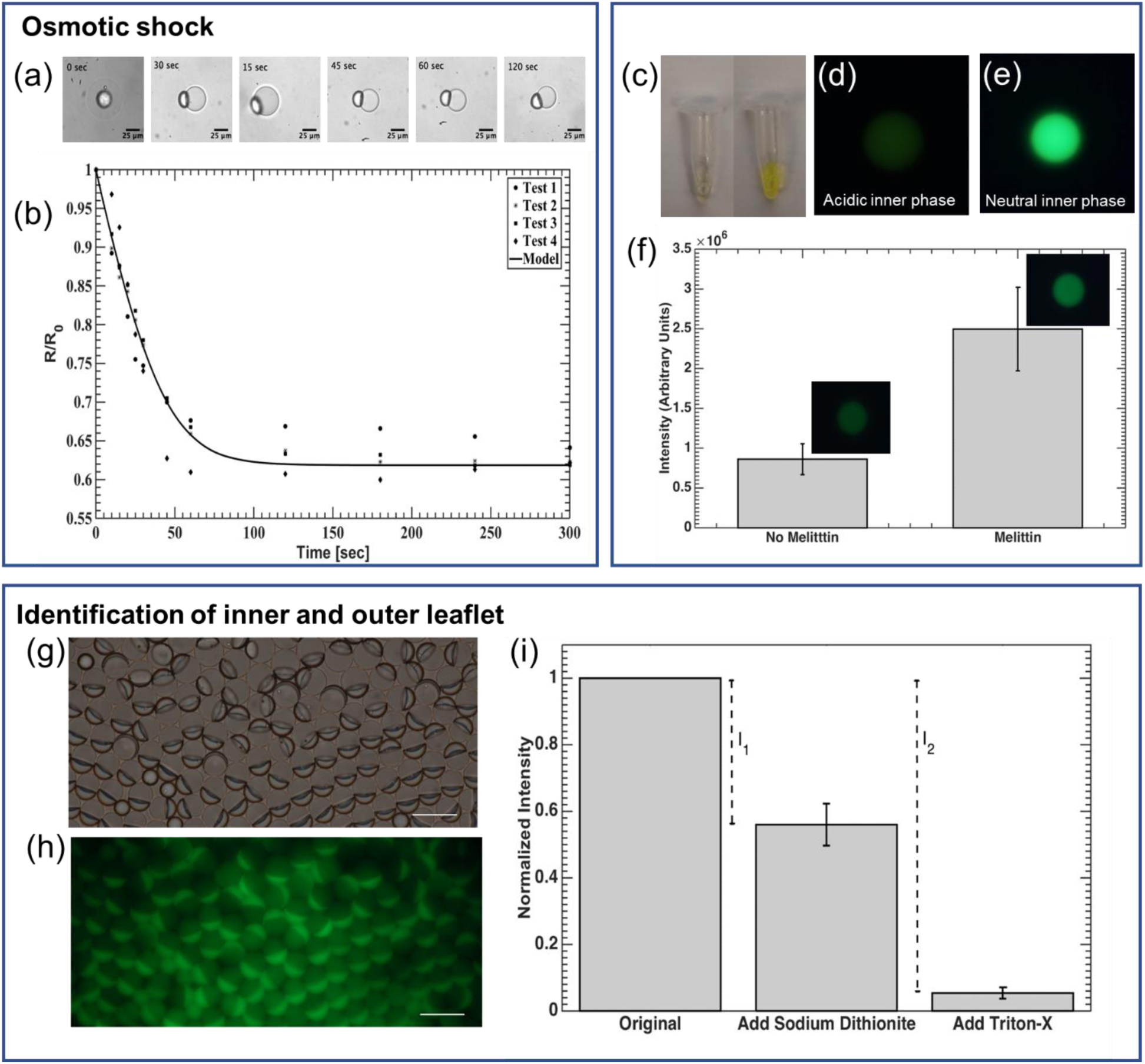
Verification of unilamellarity. (a) & (b) Permeability of SCMs measured by osmotic shock test. (a) Bright-field microscopic images of SCM shrinkage when exposed to osmotic shock. (b) SCM radius over time, normalized to the initial radius, R0, plotted when subjected to a hypertonic condition (Δc = 100 mOsm). (c-f) Insertion of the pore-forming membrane protein— melittin — into the lipid bilayer. (c) Comparison of FITC-Dextran solution color in a slightly acid NaCl solution (left, pH 4) and neutral PBS solution (right). (d) & (e) Fluorescence image of DEDs encapsulating FITC-Dextran (d) in acidic NaCl solution (pH 4) and (e) in PBS (neutral). (f) An increase in fluorescence intensity was observed for vesicles incubated with melittin for 30 minutes (p < 0.01, N = 16). (g-i) Identification of the inner and outer leaflet of SCMs. (g) Bright-field and (h) Fluorescence image of SCM population with NBD-PE incorporated in the bilayer. Excess NBD-PE can be seen in the oil caps. Scale bar = 25 μm. (i) Results of NBD-PE quenching assay. After adding sodium dithionite, the outer leaflet was quenched. I1 is the intensity contributed by the outer leaflet. After the addition of Triton-X, SCMs were destroyed, and the inner leaflet was exposed. Therefore, I_2_ is contributed by both the inner and outer leaflet. Normalized fluorescent intensity of SMC solution (λ_emssion_ = 520 nm) was used to calculate lamellarity (N = 4).

#### 2.3.2. Membrane protein insertion

Figure 3c shows that the fluorescence of FITC-Dextran was quenched in a slightly acidic solution (pH 4), and the fluorescence is bright in a neutral solution. The drastic difference in fluorescent intensity was also seen in the DEDs encapsulating FITC-Dextran in acid and neutral solution (Figure 3d& e). The fluorescent intensity remained low after DEDs were dewetted into SCMs (Figure 3f, no melittin). With the presence of melittin, the fluorescent intensity of SCMs increased significantly because the pores created by melittin are large enough for the ions in the environment solution (1X PBS) to pass through, ^[50, 51]^ and the internal pH increased as a result (Figure 3f, melittin). Considering that the pore formed by melittin can only span up to 5.56 nm, ^[50]^ this proves that the SCMs have a lipid bilayer, which is normally around 4 nm in thickness.

#### 2.3.3. Identification of the inner and outer leaflet

This method quantifies the fluorescence of the inner and outer lipids to verify the lipid distribution. The NBD on the outer leaflet of SCMs was first quenched by sodium dithionite, and the contribution of the outer leaflet was obtained (I1). Following that, Triton-X was added to break the SCMs. Therefore, the inner leaflet was quenched, and the intensity difference contributed by both the inner and outer leaflet was measured by I2. In our result, I1 / I2 = 0.46 ± 0.06, which is consistent with a unilamellar membrane with equal distribution of NBD-PE in the inner and outer leaflet (Figure 3g-i). I1 and I2 are calculated based on the intensities at the λemission = 520 nm since the intensity at this wavelength is mainly contributed by the NBD at the interface of SCMs, instead of that in the oil drops or oil caps (S 6).

### 2.4. Produce aAPC-shells, aAPC-BLMs, and oil drops from DEDs

DEDs containing 10 mol% DSPE-PEG(2000)-biotin were conjugated with anti-CD28, anti-LFA-1, and MHCII-NYESO to mimic a real cell. DEDs remained intact and less than 15% were lost after the 2-hour conjugation (S 7). The DEDs functionalized with the above-mentioned ligands are termed aAPC-shells (Figure 4). aAPC-BLMs were produced by immersing aAPC-shells in 1X PBS, the same process as dewetting DEDs into SCMs. In addition to making aAPC-BLMs, functionalized oil drops were also produced and conjugated with anti-CD28, anti-LFA-1, and MHCII-NEYSO on the surface. The number of ligands on the aAPC-BLMs is quantified by a series of fluorescent beads with a known value of molecules/bead. The number of molecules on each aAPC-BLM is 2,930 (molecules/aAPC-BLM) for anti-CD28 and anti-LFA-1, and 4,920 (molecules/aAPC-BLM) for MHCII-NYESO. Figure 5d shows the geometric intensity of the blank aAPCs (SCMs without any ligands) and the aAPC-BLMs (SCMs conjugated with all three ligands).

**Figure 4.**
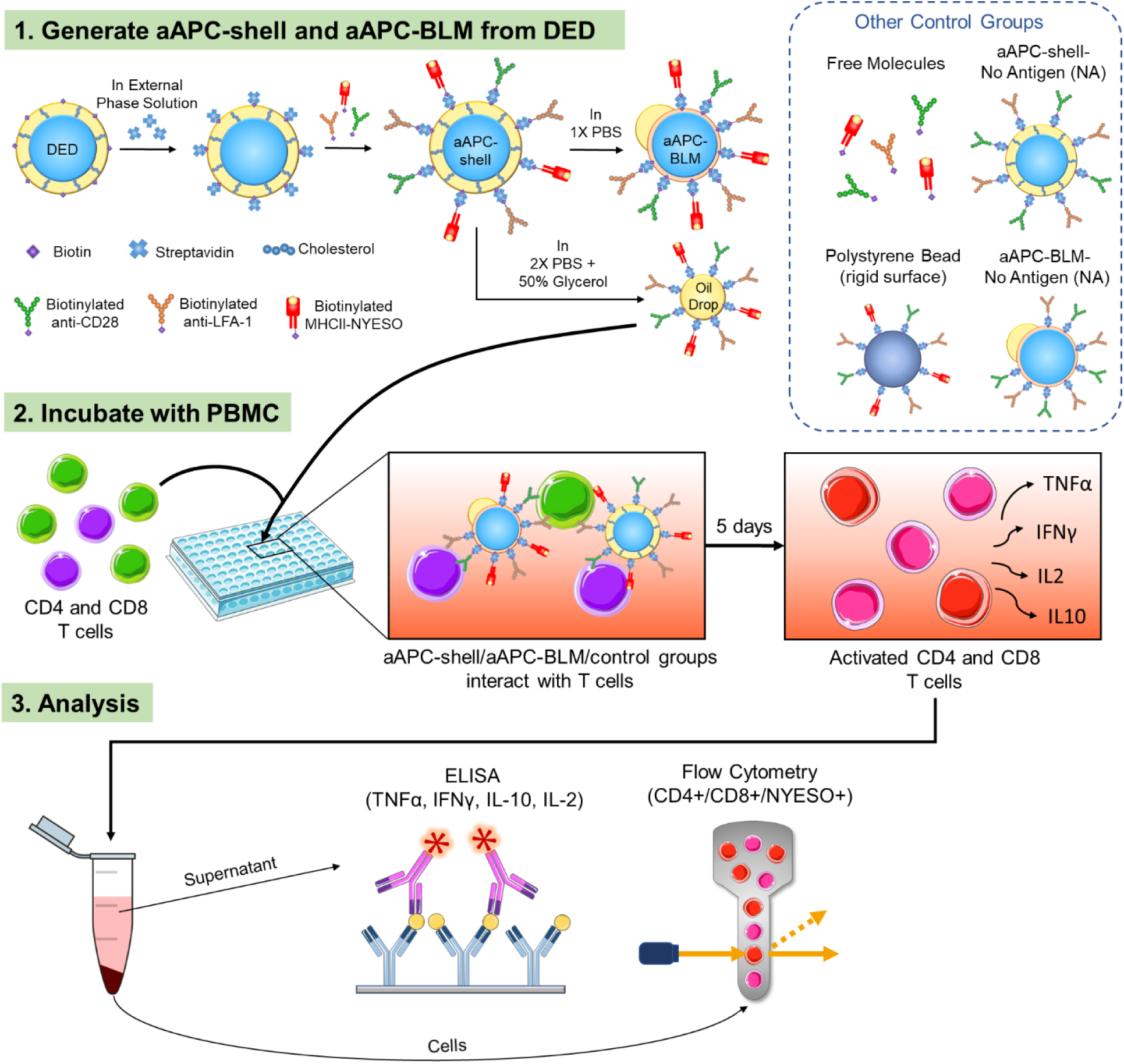
Schematic of the experimental procedure. (1) Double emulsion droplets (DEDs) are collected, conjugated first with streptavidin, and then with MHC-antigen complex and surface ligands for T cell activation. The functionalized DEDs are named aAPC-shells. The aAPC-shells are converted into aAPC-BLMs in 1X PBS or are broken into oil drops in 2X PBS + 50% Glycerol. aAPC-shells, oil drops, and other control groups are used to compare with aAPC-BLMs. (2) aAPC-BLMs are incubated with PBMC for 5 days. During the incubation, aAPC-BLMs interact with the T cells in PBMC and activate them. (3) The number of NYESO+ CD8 and CD4 T cells are quantified by flow cytometry and the cytokines (TNFα, IFNγ, IL-10, and IL-2) secreted by the cells are analyzed by ELISA.

**Figure 5.**
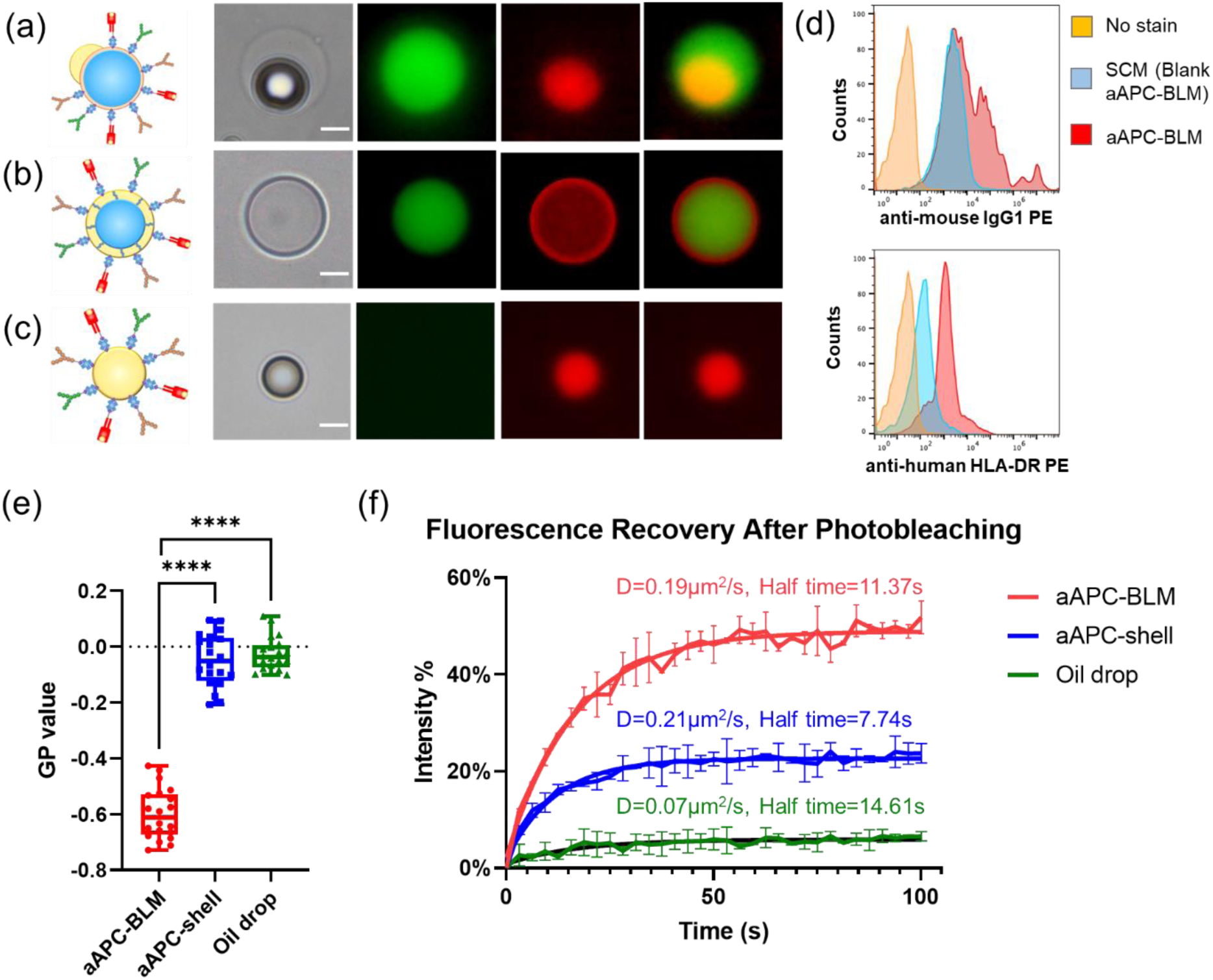
Fluidity analysis of aAPC-BLM, aAPC-shell, and oil drop. Images of (a) aAPC-BLM, (b) aAPC-shell, and (c) functionalized oil drop. The green color is the FITC inner phase, and the red color is the rhodamine-conjugated lipid. (a) In aAPC-shells, the inner phase is well encapsulated and evenly surround by the lipids. The lipids are shown as red rings. (b) In aAPC-BLMs, the inner phase is surrounded by lipid bilayers, which is too thin to be visualized. The oil cap is shown as a red dot that is not completely spherical. (c) Oil drops can be identified by the red sphere without the inner phase. Scale bar = 10 µm. (d) The intensity of the anti-mouse IgG1 PE and anti-human HLA-DR PE. (e) The fluidity of lipid bilayer, oil sell, and oil drop measured by LAURDAN generalized polarization (GP) value. A lower GP value refers to higher fluidity. N = 20 for each condition. (f) Fluorescence recovery after photobleaching (FRAP). The mobile fraction of aAPC-BLM, aAPC-shell, and oil drops are 48.69%, 22.57%, and 5.84%, respectively.

Since some (< 15%) aAPC-shell and aAPC-BLMs inevitably popped during the conjugation process, it is worth investigating whether the functionalized oil drops alone would activate T cells. Functionalized oil drops were made by breaking the DEDs with a high-osmolarity solution (2X PBS + 50% glycerol). The images of aAPC-shell, aAPC-BLM, and functionalized oil drop are shown in Figure 5a-c. TEM images of SCM and oil drop also show the morphology of the lipid-bilayer film, and the fatty acid/lipid cluster that is consistent with our observation under the fluorescence microscope (S 8).

Several control groups were added to verify the effect of membrane fluidity and size (Figure 4). Antibody-coated polystyrene beads with a diameter of 16.5 µm were used as a rigid-surface control (S 9). Anti-CD28, anti-LFA-1, and MHCII-NYESO molecules were suspended in the solution to test whether T cells can be activated without the mechanical force induced by the micron-sized droplet. aAPC-shells and aAPC-BLMs without MHCII-NYESO presented on the surface (aAPC-shell-NA and aAPC-BLM-NA — NA denoting no antigens) were used to know the baseline of T cell response when exposed to aAPC-shells and aAPC-BLMs. aAPC-BLMs conjugated with mycobacterium tuberculosis (termed aAPC-BLM-TB) were tested to identify whether the activation is antigen-specific. The NYESO peptide 157-170 (SLLMWITQCFLPVF) displayed on the surface is a CD4 T cell epitope. However, the amino acids 157-165 (SLLMWITQC) also represented the NYESO CD8 T cell epitope. This peptide was specifically chosen to determine if our aAPCs will display MHC restriction. If MHC restriction was present, then when the NYESO peptide is coupled to MHCII, it should not activate CD8 T cells. To investigate this, both CD4 and CD8 T cells were analyzed (section 2.8).

### 2.5. aAPC-BLM has a higher mobile fraction than aAPC-shell and oil drops

The fluidity of aAPC-BLM, aAPC-shell, and oil drop was measured by incorporating LAURDAN in the oil phase. The long lipophilic tail allows it to settle on the lipid-bilayer membrane of aAPC-BLM or the oil-water interface of aAPC-shell and oil drop. LAURDAN is a fluorescent probe that is sensitive to the local lipid packing. ^[52–55]^ When the lipid-bilayer membrane is in the gel phase or liquid-ordered phase (lo), water molecules are mostly excluded from the closely packed lipids. When the membrane is in the liquid-disordered phase (ld), more water molecules can be found in the proximity of LAURDAN. As LAURDAN is excited, the energy is used to reorient the water molecules, causing the dipolar relaxation of water. The emission spectrum experiences a red-shift since a part of the energy is consumed in dipolar relaxation. On the other hand, in the lipid-ordered phase (lo), minimal dipolar relaxation can occur, therefore the emission spectrum has a shorter wavelength.

The generalized polarization (GP) value was calculated based on the normalization of the emission intensity of the liquid-ordered phase minus that of the liquid-disordered phase (S10). Therefore, a lower GP value represents a higher GP value. The GP values of aAPC-BLM, aAPC-shell, and oil drop are -0.597 ± 0.089, -0.048 ± 0.092, and -0.026 ± 0.059, respectively. This shows that aAPC-BLM is much more fluid than aAPC-shell and the oil drop (Figure 5e).

In addition to LAURDAN, which is a good detector for local fluidity, fluorescence recovery after photobleaching (FRAP) was performed to investigate the fluidity on a larger scale (Figure 5f). aAPC-BLM, aAPC-shell, and oil drops were generated using rhodamine-labeled lipids. The region of interest (ROI) was photobleached by a laser beam with strong intensity. After photobleaching, rhodamine-labeled lipids in the unbleached area diffused into the photobleached area, thus the fluorescent intensity recovered over time.^[56]^ The FRAP analysis shows that the oil drop has a low diffusion coefficient (D) of 0.07 µm2 s-1 and a long half-life of 14.6 s. The aAPC-shell has a slightly higher diffusion coefficient (D = 0.21 µm2 s-1) and a shorter half-life (7.74 s) than aAPC-BLM (D = 0.19 µm2/s, half-life = 11.37s). Based on the Stokes-Einstein equation 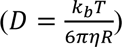, ^[57]^ diffusivity is inversely proportional to the viscosity (η) of the liquid. The significantly lower diffusion coefficient and higher half-time of the oil drop are attributed to the viscous oleic acid. In a 2D thin film, the diffusion coefficient, also known as diffusivity, is calculated by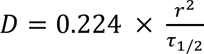. ^[58–60]^

It is important to note that the diffusivity is dependent on the properties of the materials, therefore the lipids in the bilayer and the oil shell are not fully comparable. While the half-life of the aAPC-shell is shorter than aAPC-BLM, the diffusivities are close, and more importantly, the mobile fraction of aAPC-BLM (48.68%) is over two times higher than that of aAPC-shell (22.57%). The mobile fraction reflects the lateral mobility of the lipids. According to Mossman et al., ^[61]^ the size of an immune synapse is 5-6 µm in diameter, and the photo-bleached ROI in this experiment is a circle of 5-7 µm diameter, suggesting the FRAP result could reflect the diffusivity in an actual immune synapse. The higher mobile fraction indicates that more ligands are able to diffuse into the area where the immune synapse forms.

### 2.6. aAPC-BLMs can induce significant higher amount of cytokines secretion than aAPC-shells in PBMC

The number of aAPC-BLMs for T cell activation was optimized to minimize the toxicity while maintaining the efficacy (S11). In our study, the number of T cells was fixed at 50,000/well, and aAPCs were tested from 1,500 to 6,000 per well. More distinguishable toxicity was found when aAPCs reached 6,000/well. Based on our findings, 3,000 aAPCs/well was sufficient to activate T cells while preventing cell death. Oleic acid is the main source of toxicity. Studies have reported that oleic acid promoted apoptosis and necrosis in Jurkat and human lymphocytes, and the viability of cells dropped at 200 µM oleic acid. ^[62, 63]^ 3,000 aAPCs/well corresponds 38 µM of oleic acid, which is well below the reported toxic concentration.

With the stimulation of aAPC-BLMs, the secretion of IFNγ, TNFα, IL-10, and IL-2 in PBMC increased by 28, 7, 17, and 9.6-folds, respectively (Figure 6a&b). IFNγ, TNFα, and IL-2 are secreted when the T cells are activated. IL-2 promotes the proliferation of activated T cells. IL-10 provides information on the immune-regulation of T cells. When the cells are constantly signaled to be activated, IL-10 is secreted to balance the activation. Over-activation may cause exhaustion. IL-10 secretion could keep T cells viable and cytotoxic (Figure 7e) after 5 days of co-culturing with aAPC-BLM.

**Figure 6.**
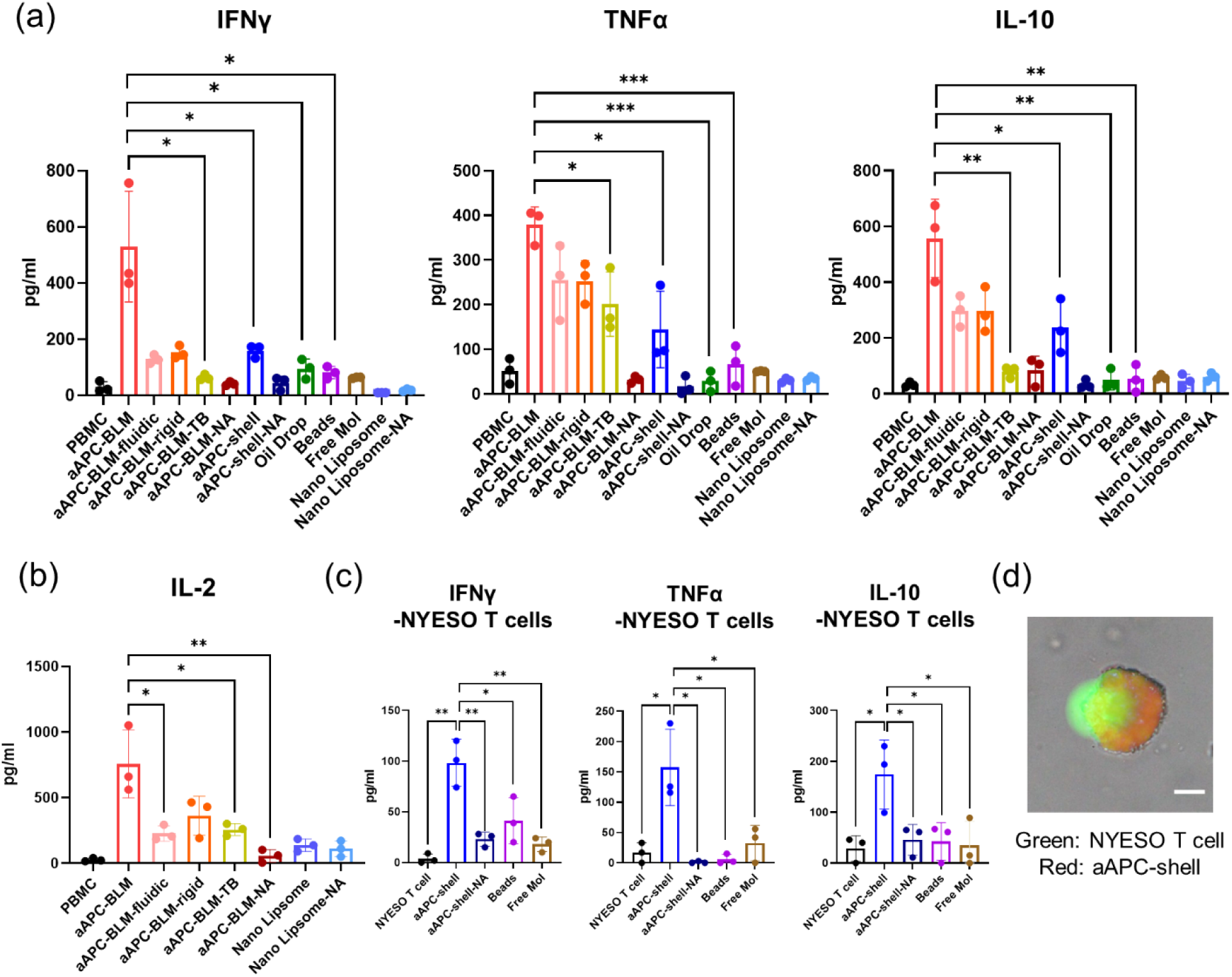
Analysis of (a) IFNγ, TNFα, IL-10, and (b) IL-2 in the supernatant after co-culturing the test articles with PBMC for 5 days. (c) Cytokine secretion of NYESO-specific CD8 T cells after 5 days of incubation with aAPC-shell or control groups. (d) CFSE-stained NYESO-specific T cell interacting with aAPC-shell (prepared with 18:1 Liss Rhodamine-PE). T cell maximized its contact area with the aAPC-shell, indicating the immune synapse was formed. Scale bar = 10 µm.

**Figure 7.**
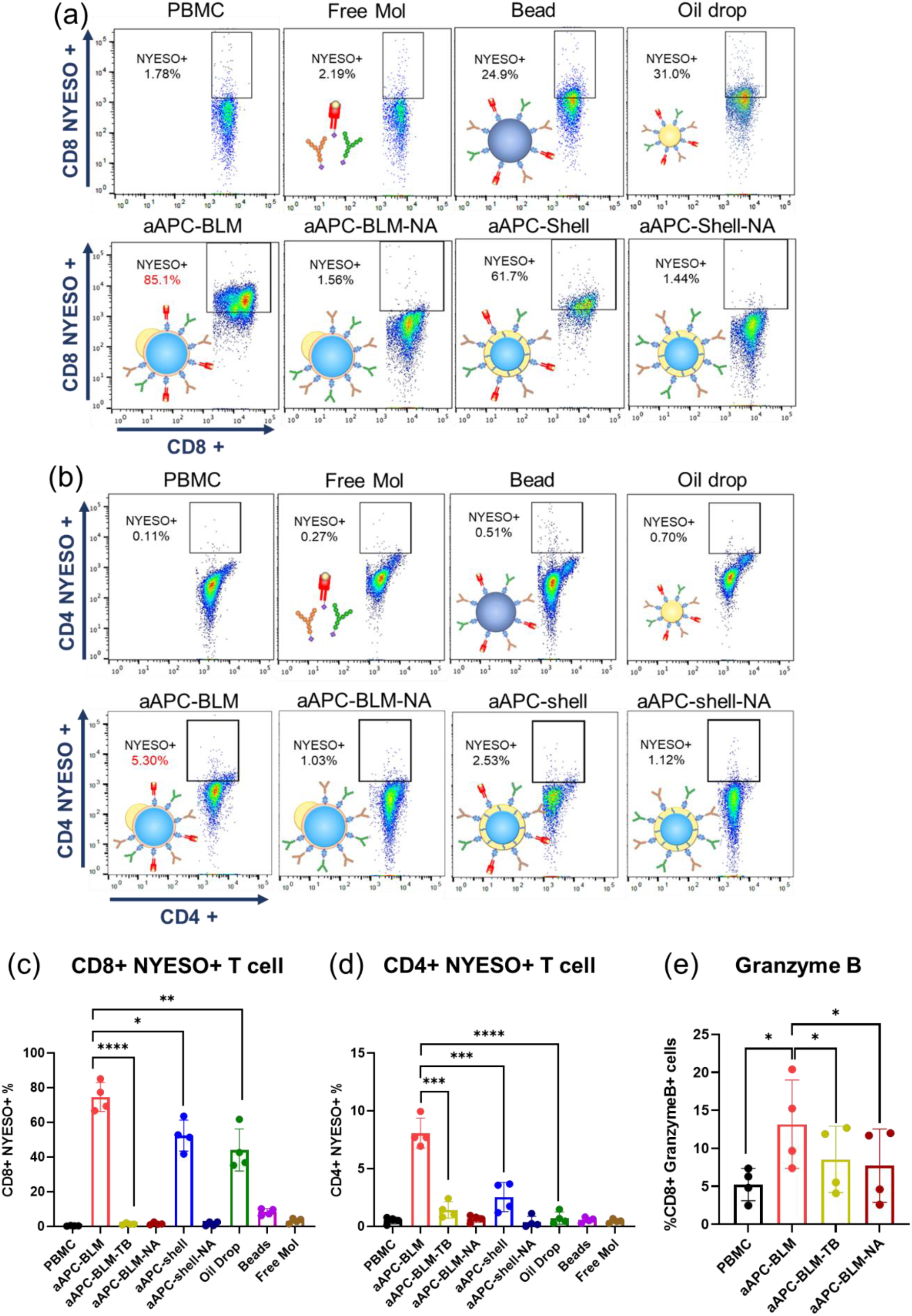
Analysis of NYESO+ CD4 and CD8 T cells in PBMC after incubating with aAPC-BLMs for 5 days. NYESO tetramer staining of (a) CD8+ T cells and (b) CD4+ T cells in PBMC. (c) Results for (a). (d) Results for (b). (e) Intracellular staining of granzyme B.

aAPC-BLMs induced significantly higher cytokine secretion compared to aAPC-shell and functionalized oil drops, which suggests that the fluidic membrane is crucial to T cell activation. To further investigate how fluidity affects the T cell response, aAPC-BLMs with different fluidities were generated by tuning the cholesterol concentration. The result shows that the aAPC-BLM induced a significantly higher cytokine secretion than aAPC-BLM-fluidic and aAPC-BLM rigid. Although it was hypothesized that the higher fluidity would induce a stronger T cell response, there was no notable difference in the cytokine secretion induced by aAPC-BLM-fluidic and aAPC-BLM-rigid. It was also possible that the aAPC-BLM has the proper fluidity that elicit the most effective interaction with cells, but it should be stressed that aAPC-BLM-fluidic and the aAPC-BLM-rigid are less stable than aAPC-BLM. They easily popped into oil drops (S 12), therefore, the effect of the fluidic BLM was compromised.

aAPC-shells induced the secretion of IFNγ and IL-10 in PBMC by 4 and 7-folds, respectively, compared to non-stimulated PBMC. However, they were unable to induce a significant increase in TNFα, whether compared with non-stimulated PBMC or other control groups. Functionalized oil drop, beads, aAPC-shell-NA, aAPC-BLM-NA, and free molecules were not able to stimulate T cells. aAPC-BLM-TBs induced a low level of TNFα and IL-2 compared to aAPC-BLM (Figure 6a&b) and could not expand NYESO+ T cells (Figure 7b&d). The statistical analysis of the comparisons is presented in S 13. The nanoliposomes (125.5 nm in diameter, S 14) did not induce meaningful cytokine secretion, even when administered 10 times more than the aAPC-BLMs. This shows that having a fluidic surface alone is not sufficient for T cell activation.

### 2.7. aAPC-shells induce significant cytokine secretion in NYESO-specific T cells

As aAPC-shells did not show significant TNFα secretion in PBMC, it is of interest to know whether they can induce cytokine secretion in antigen-specific CD8 T cells. Figure 6c shows that aAPC-shells were able to induce the secretion of IFNγ (27-fold), TNFα (9-fold), and IL-10 (6-fold) in NYESO-specific CD8 T cells after 5 days of incubation. This suggests that the aAPC-shells can be used as boosters or therapeutics for cancer patients.

The induction of cytokine secretion can be evidenced by the findings that aAPC-shells could interact with T cells. aAPC-shells were able to associate with NYESO-specific T cells within 30 minutes, and within 2 hours, a single aAPC-shells could bind with multiple T cells (S 15a). The immune-synaptic association between T cells and aAPC-shells was also observed in 2 hours, as the T cell were expanding at the contact junction and spreading over the aAPC-shells (Figure 6d). This is in line with the studies that immune synapses can form in 5-10 minutes upon TCR-MHC-peptide conjugation and mature in 30 minutes. ^[64, 65]^ After incubation for 18 hours, the interaction between a single aAPC-shell and multiple T cells still existed (S 15a). Our finding is well in line with the studies by Von Andrian et al. that stable, longer-lasting synapses formed after 12 hours of APC-T cell interaction. ^[66, 67]^ Comparatively, much fewer beads were found to be interacting with the T cells throughout 18-hour of observation (S 15a).

To confirm whether the contact was a tight immune-synaptic interaction or was merely the result of T cells and aAPCs/beads settling in proximity, the solution was pipetted to break the loose contact, and 10 µL was transferred into a countess slide for observation. It was shown that multiple T cells were still engaging with aAPC-shells after vigorous pipetting (S 15b). However, no bead-T cell association was found after the pipetting, suggesting that the beads cannot form a sturdy immune synapse with T cells as aAPC-shells do.

### 2.8. aAPC-BLMs expand significantly higher NYESO-specific CD4 and CD8 T cells than aAPC-shells in PBMC

The proliferation of antigen-specific T cells is a key indicator of T cell activation. aAPC-BLMs are co-cultured with PBMC for 5 days with a ratio of 1:17 (aAPC:cells), which is much lower than the DynabeadsTM where a ratio of 1:1 is required. Despite the ratio, aAPC-BLMs had a highly effective activation of both NYESO-specific CD8 and CD4 T cells in PBMC, where the NYESO-specific CD8 T cells were expanded from 0.32% to 74.6% (233-fold), and the NYESO-specific CD4 T cells from 0.5% to 8.1% (16-fold) (Figure 7, S 16). The result shows that the activation is not MHC-restricted. In comparison, aAPC-shell expanded the CD8 T cells by 164-fold and CD4 T cells by 5-fold. Considering both T cell expansion (Figure 7a-d) and cytokine secretion result (Figure 6a&b), it was found that aAPC-BLMs were more effective than aAPC-shells. It is conjectured that while aAPC-shells have a fluid oil shell surface, they do not have the lipid bilayer as real cells do. aAPC-BLMs, on the other hand, could recapitulate the lipid bilayer fluidity of a real cell, thus contributing to higher efficacy. While oil drops could induce 137-fold expansion of CD8 T cells, they are ineffective in inducing CD4 T cell expansion. The beads were ineffective in activating either CD8 or CD4 T cells. The overall underperformance of the beads is likely due to the rigid surface restricting the formation of the immune synapse. Free molecules also did not result in T cell expansion. This is conjectured to be due to the absence of mechanical force that is present in cell-cell synapses and critical for the initiation of signal transduction. aAPC-shell-NAs, aAPC-BLM-NAs, and aAPC-BLM-TB did not stimulate the expansion of NYESO-specific T cells, which proves that the activation is antigen-specific. The cells are not only antigen-specific but also cytotoxic, as evidenced by the upregulated secretion of granzyme B (Figure 7e).

Compared to the aAPCs reported in other studies, aAPC-BLMs have higher efficacy in many aspects. First of all, priming and boosting are not required. Priming the T cells with peptide-pulsed DCs before aAPC stimulation or restimulating the cells multiple times are required in many studies.^[18]^ ^[68]^ Rudolf et al. presented an average of 5-fold increase in antigen-specific T cells after 3-4 times of repeated stimulation, ^[69]^ while our study demonstrated more than two-hundred-fold increase for CD8 T cells without any boosting. Although some studies do not require repeated stimulation, they do require a long incubation time (28 days - 9 weeks) to acquire enough cytotoxic T cells, ^[70, 71]^ whereas 5 days of incubation was sufficient in our case. Secondly, both aAPC-BLMs and aAPC-shells are effective without incorporating cytokines, adjuvants, ^[72]^ or other surface molecules. Some studies needed to encapsulate cytokines and chemokines, such as IL-2, IL-7, IL-15, IL-21, and CCL-21, in the particles or add them into the cell culture medium. ^[73]^ Others conjugate anti-4-1BB to enhance the amplification of the activated T cells, ^[69, 74]^ or use anti-PD-1 to prevent programmed cell death. ^[75]^ In our study, aAPC-BLMs circumvent the need for any stimulants. Thirdly, the aAPC-to-T cell ratio for effective activation is lower than that in most studies. The ratio of aAPC:T cell ranges from 1:1 to 1:8 in most studies, ^[74, 76]^ while a ratio of 1:17 is enough to elicit an immune response in this study. Last but not the least, we demonstrated the expansion of antigen-specific T cells, which is more challenging than polyclonal expansion, ^[19, 74]^ since the presence of antigen-specific T cells is very rare in healthy donors. The antigen-specific T cells have been reported to increase by 2-40 folds in general, with the addition of aAPCs along with other stimulatory molecules. ^[77]^ In comparison, aAPC-BLMs can activate T cells by up to 233 folds in a simpler manner (without boosting, without stimulant, with short incubation time, with a low aAPC-to-T cell ratio). Finally, the aAPC-BLM platform has the potential to be applied to a variety of antigens, such as tetanus (S 17) and treatments, such as inducing insulin secretion in pancreatic β cells.^[34]^ In future studies, the efficacy can potentially be further enhanced by encapsulating some signaling molecules and adjusting the stiffness of aAPC-BLMs.

## 3. Conclusions

This research presents a method to generate monodisperse lipid bilayer aAPCs (aAPC-BLMs) that could successfully activate the rare NYESO-specific T cells in PBMC, showing up to 233-fold of CD8T cell expansion and 28-fold of cytokine secretion. In addition to the aAPC-BLMs, aAPCs with a thin oil shell (aAPC-shell) were produced to investigate the effect of lipid bilayer. While aAPC-shell was not as effective as aAPC-BLM in PBMC T cell activation, it could form an immune synapse with formerly primed NYESO-specific T cells and elicit distinctive cytokine responses, suggesting its promise of being applied as boosters. The study also proves that size and membrane fluidity are crucial factors to T cell activation. The cell-like lipid bilayer membrane of aAPC-BLMs contributes to their greater efficacy over aAPC-shells.

The aAPC-BLMs presented in this study have many advantages over other current aAPCs. First of all, the T cells can be activated with a low aAPC-to-T cell ratio and a relatively short incubation time (5 days). The aAPCs have high stability, controllable size, and tunable surface fluidity. This is a flexible and versatile tool to study the optimized physiological properties of aAPC. For instance, the fluidity can be tuned by changing the lipid composition, the stiffness can be adjusted by encapsulating different concentration of hydrogels, and the density of the surface molecules can be varied by incorporating different concentration of biotin. This easy-to-perform method will greatly reduce the time, cost, and labor for culturing natural APCs. Furthermore, this offers a way to produce aAPCs with high quality, consistency, scalability, and can be easily preserved. The combined advantages make this method and the as-prepared aAPC-BLMs exceptionally promising to improve the status quo of ACT immunotherapy. To the best of our knowledge, no other studies have successfully activated T cells using cell-sized lipid-bilayer vesicles as aAPCs. The aAPC-BLMs exhibited great potency without the need for priming, restimulation, or the use of stimulatory molecules. aAPC-BLMs could be administered at a lower concentration while still maintaining a higher efficacy than most current aAPCs. The low-cost, high stability, scalability, and short processing time make competitive as an off-the shelf aAPC product. This method allows for simple modification of the ligand combination, ligand density, fluidity, and other properties to create personalized T cell activation treatments. Further studies could build on this platform to unveil the fundamental contributing factors of T cell activation. The future impact is that patients will be able to customize the aAPCs for their personalized treatments and strengthen the field of adoptive cell transfer for the foreseeable future.

## 4. Experimental Methods

### 4.1. Materials

18:1 (Δ9-Cis) PC (DOPC) (1,2-dioleoyl-sn-glycero-3-phosphocholine), 16:0 PC (DPPC) (1,2-dipalmitoyl-sn-glycero-3-phosphocholine), DSPE-PEG(2000)-biotin (1,2-distearoyl-sn-glycero-3-phosphoethanolamine-N-^[biotinyl(polyethylene glycol)-2000]^ (ammonium salt)), and 16:0 Liss Rhod PE (1,2-dipalmitoyl-sn-glycero-3-phosphoethanolamine-N-(lissamine rhodamine B sulfonyl) (ammonium salt)), DOPG (1,2-dioleoyl-sn-glycero-3-phospho-(1’-rac-glycerol)), 16:0 NBD-PE (1,2-dipalmitoyl-sn-glycero-3-phosphoethanolamine-N-(7-nitro-2-1,3-benzoxadiazol-4-yl) (ammonium salt)) were purchased from Avanti Polar Lipids. Oleic acid was purchased from Tokyo Chemical Industry (TCI America). Cholesterol and polyvinyl alcohol (PVA, MW 30,000– 70,000, 87%–90% hydrolyzed) were purchased from Sigma Aldrich. Pluronic™ F-68 Non-ionic Surfactant (100X). Sucrose, isopropanol alcohol (IPA), Phosphate-Buffered Saline (PBS) and RPMI 1640 medium were purchased from Thermo Fisher. Glycerol was purchased from EMD Biosciences. Biotin anti-human CD28 antibody (Cat#302904), biotin anti-human CD11a/CD18 (LFA-1) antibody (Cat#363424), and FITC anti-mouse IgG Antibody were purchased from BioLegend. Streptavidin eFluor 570 was purchased from eBiosciecne. NYESO-1-MHCII biotinylated monomer (DPB1*04:01/DPA1*01:03 human CTAG1 157-170 SLLMWITQCFLPVF) and mycobacterium tuberculosis biotinylated monomer (DRB1*03:01 Mtb groEL 1-13 MAKTIAYDEEARR) were provided by NIH Tetramer Core Facility. The Polydimethylsiloxane (PDMS) and its curing agent was purchased from Dow Corning Sylgard 184 Silicone Elastomer Kit. SU8-10 and SU8-2010 were purchased from MicroChem. Biotin-coated polystyrene beads (Cat#TPX-150-5) were purchased from Spherotech.

### 4.2. NYESO-specific T cells, peripheral blood mononuclear cells, ELISA kit, and reagents for tetramer staining

NYESO-specific CD8 T cells were obtained from Cellero (Lowell, MA). CD4 FITC (clone A161A1) and CD8 PerCP (Clone SK1) were purchased from Biolegend (San Diego, CA). NYESO APC tetramer and CD4 NYESO PE tetramer were offered by NIH Tetramer Core Facility at Emory University, Atlanta. IL-10, IFNγ and TNFα ELISA kit were purchased from BD Biosciences (San Jose, CA). Peripheral blood mononuclear cells (PBMCs) were isolated from blood of healthy subjects by Ficoll-hypaque density gradient centrifugation. Blood was collected under the protocol approved by Human Subject Committee of the Institution Review Board of the University of California, Irvine.

### 4.3. Channel design

The water-in-oil-in-water (w/o/w) DEDs were generated in a single-step manner with a flow-focusing channel (Figure 2) where the internal aqueous phase (250 mM sucrose with 1% F68), the middle oil phase (lipids and cholesterol dissolved in oleic acid), and the external aqueous phase (15% glycerol and 6% F68 with 125 mM NaCl) converged at a 15 µm orifice. An array of 10 µm filters was designed to trap the debris at all three inlets, and a winding long channel was used to regulate the flow. Compared to the two-step droplet generation ^[78]^, single-step droplet generation is able to produce DEDs with an oil shell that is significantly thinner. The benefits of having a thin oil shell include increasing the stability for long-term storage and reducing the time for oil removal to form lipid-bilayer ^[79]^, which is beneficial to cell-mimicking. Due to these reasons, single-step design was adopted.

### 4.4. Fabrication of the silicon wafer mold and PDMS channel

The microfluidic channel was made by casting PDMS onto a silicon wafer mold fabricated by photolithography (S 1). The channel geometry was designed using AutoCAD and the photomask was printed through CAD/Art Services. The wafer was fabricated in two layers, with the first layer being 30 µm and the second layer being 10 µm. The second layer was designed as a step to reduce the channel height, located before the internal phase channel merged with the oil phase channel to avoid the wetting of internal aqueous phase. The first layer (30 µm) was produced by coating the photoresist SU8-10 on a 4-inch silicon wafer. After photo-crosslinking the pattern and removing the residual photoresist, the wafer was baked at 220 ℃ for 10 minutes for annealing and solvent evaporation. Following that, the second layer of photoresist (SU8-2010) was spin-coated onto the same wafer and the mask for the second layer was aligned using MA56 Mask Aligner. Both layers were fabricated by following the MicroChem protocol. To prevent the adhesion of PDMS to the wafer, the patterned wafer was spin-coated with the mixture of PTFE (polytetrafluoroethylene) and FC40 (volume ratio = 1: 5) at 3000 rpm for 30 seconds. After the spin coating, the wafer was heated at 120 °C for 20 minutes to ensure the complete evaporation of liquids. For replicating the pattern, PDMS was mixed with the curing agent at a weight ratio of 10:1. The mixture was poured onto the wafer, degassed for 30 minutes, cured at 65 °C overnight, and finally the PDMS would be ready for assembly. The single-layer DED trapping array was fabricated at 30 µm height following the same procedures.

### 4.5. Device assembly and PVA treatment

The final microfluidic device was made by assembling the PDMS channel and the PDMS-coated glass slide (75 x 51 mm, 1.2 mm thick, purchased from VWR). The reason for using a PDMS-coated glass slide instead of the glass slide itself is because PDMS decreases the wettability of internal phase, plus it makes the channel property uniform on all sides. The PDMS-glass slide was prepared by spin-coating the glass slide with the mixture of PDMS and curing agent in two steps (1st step: 3000 rpm, 30 s, 300 rpm/s; 2nd step: 2100 rpm, 50 s, 100 rpm/s). After overnight curing at 65 °C, PMDS-coated slide was ready for use. On the other hand, the cured PDMS channel was cut out and peeled off from the wafer, and the inlets and outlet were punctured with a 1.5 mm biopsy punch (Integra™ Miltex®).

To assemble the punctured PDMS channel and PDMS-coated slide, both compartments were treated with air plasma (Harrick Scientific) at 300 torr for 2 minutes. Next, the compartments were tightly sealed, and the external-phase channel was treated with PVA solution to ensure the hydrophilicity of the selected channel. The PVA solution was prepared by dissolving 1 wt% of PVA in DI water. The dissolving of PVA was assisted by heating at 95 ℃ while mixing with a magnetic stir bar at 200 rpm. The solution was cooled down for one day, diluted to 0.4% and 0.1%, and filtered with 0.2 µm filters (cellulose acetate membrane, VWR).

The PVA treatment was done by applying vacuum at the outlet while adding 2 µl of 0.4% PVA to the inlet of the external-phase channel. After the 0.4% PVA was vacuumed out, 2 µl of 0.1% PVA was added to the inlet to wash away PVA crystals that was formed during the 0.4% PVA treatment. After the 0.1% PVA was added, the vacuum was applied for 5 minutes to ensure the solution was completely removed. The PVA-treated device was heated at 120 ℃ overnight so that the non-treated area could return to hydrophobicity.

### 4.6. Double emulsion droplet generation and preservation

The water-in-oil-in-water (w/o/w) double emulsion droplets (DEDs) were generated using a flow-focusing microfluidic device. The internal aqueous phase was composed of 250 mM sucrose with 1% F68 and 2 mg ml^-1^ 70 kDa dextran-FITC. The middle oil phase was prepared by dissolving DOPC (7.5 mg ml^-1^), DPPC (2.5 mg ml^-1^), cholesterol (5 mg ml^-1^), 0.05 mol% 16:0 Liss Rhod PE, and 5 or 10 mol% DSPE-PEG(2000)-biotin into oleic acid. The external aqueous phase was composed of 15% glycerol and 6% Pluronic F68 with the addition of 125 mM NaCl for osmolarity balance. The experimental setup was described in our previous publication.^[34]^ In brief, a LabView-controlled pressure pump was used to push the fluids through Tygon tubing (Cole Parmer, ID = 0.2 in, OD = 0.6 in) into the channels, and a tubing was inserted at the outlet to collect the DEDs.

The pressure for internal, middle and external phase were set to be 0.8 psi, 1.6 psi, and 3.6 psi, respectively. The pressure could vary by ± 0.3 psi each time due to the inevitable minor differences among each fabrication. Droplet generation was monitored and recorded using Phantom High-Speed Camera (V310 Phantom, Vision Research). The DEDs were preserved in the external phase solution for long-term storage. The DEDs used for the antibody conjugation are prepared in a week before the experiment.

### 4.7. Trapping array design and fabrication

Each trap was designed to have a half-circular shape with a gap at the center. The opening is 60 µm and the gap is 10 µm. The mold was made by photolithography on a silicon wafer, as described in the previous section. The height was made to be 40 µm. PDMS was casted onto the mold to replicate the structure. The inlet was punched with a 4 mm-diameter hole as a reservoir, and the outlet was punched with a 1.5 mm-diameter hole. The PDMS slab with the array pattern and a clean glass slide were plasma bonded right before the experiment, so that the channel would be hydrophilic while loading the droplets. A Tygon tubing (Cole Parmer, ID = 0.2 in, OD = 0.6 in) connected to a syringe pump was inserted at the outlet. A flow rate of 10 µl min-1 in withdrawal mode was applied at the beginning. After the droplets flew into the channel, the flow rate was decreased to less than 3 µl min-1. For overnight observation, fill the inlet reservoir with RPMI, turn off the syringe pump, but leave the tubing on the chip, so that the negative pressure would remain.

### 4.8. Verification of unilamellarity

4.8.1. Osmotic shock

The osmotic shock test was conducted by placing SCMs in a hypertonic solution that is 100 mOsm higher than the inner core of SCMs. The size of SCMs was recorded till equilibrium was reached. Since the volume of the external solution is much larger than the droplets, it is assumed that the concentration of the external solution (cext) remains the same. Therefore, Equation (1) can be written as Equation (2).

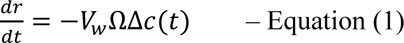

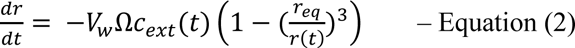

*r* = *radius*; *V_w_* = *molar volume of water*; Ω = *membrane permeability*

Δ*c*(*t*) = *the difference in solution concentration between the interior of the SCM and the external solution*

By measuring the SCM radius over time, the permeability of the membrane was calculated based on Equation (2).

#### 4.8.2. Membrane protein insertion

The bilayer was further confirmed by incorporating the transmembrane protein— melittin— into the SCMs. Melittin is a pore-forming peptide that self-assembles on the surface and inserts through the membrane. During droplet generation, 100 µM of FITC-Dextran in a slightly acid (pH 4) NaCl solution was encapsulated into DEDs. Fluorescein is a pH-sensitive dye that shows weak fluorescence under acidic conditions but has bright fluorescence at a neutral pH. A 7:3 molar ratio of DOPC : DOPG was used to improve melittin incorporation into the membrane ^[50]^. DEDs were dewetted into SCMs in 1X PBS. After that, melittin was added into the SCM solution to reach an effective melittin concentration of 4.5 μM. SCMs were incubated in the melittin-containing PBS solution for 30 minutes before observation.

#### 4.8.3. Identification of the inner and outer leaflet

A fluorescence-quenching assay is used to measure the fraction of outer leaflet ^[80–82]^. If the fluorescent intensity is reduced by half after the fluorescence quenching, the lipids are in the form of a bilayer. DEDs including 5 mol% of 16:0 NBD-PE were first generated and converted into SCMs. The contribution of the outer leaflet was calculated by quenching the fluorescence of NBD at the surface of SCMs. 10 vol% of 1M sodium dithionite in Tris buffer (pH 10, freshly prepared) was added to the solution to quench the fluorescence. As sodium dithionite cannot diffuse into the membrane, it only quenches the fluorescent intensity of the outer leaflet. The intensity difference before and after quenching was denoted by I1, which is the contribution of the outer leaflet. Following that, 1.25 vol% of 20 % Triton-X was added to identify the contribution of the inner leaflet. The SCMs were destroyed by Triton-X, consequently, the inner leaflet was exposed to the sodium dithionite-containing solution and was quenched. The difference between the initial and final intensity was denoted as I2. 400 μl of this solution containing the SCMs was placed in a cuvette, and the emission spectrum (480 nm – 600 nm) was measured using a SPEX Fluorolog 1680 0.22 m Dual Spectrometer, with the excitation wavelength set to 465 nm. Intensities at the emission wavelength of 520 nm (λ_emission_ = 520 nm) was analyzed. All the intensities are normalized based on the initial intensity. The value of I1 / I2 is equal to 50 % if the lipids are unilamellar. ^[80–83]^

### 4.9. Generation of aAPC-shell, aAPC-BLM, and oil drop

In the body, T cells are activated by APCs via the association of TCR to MHCII-peptide, CD28 costimulatory molecules to CD80/CD86, and LFA-1 adhesion molecules to ICAM-1, forming an immune synapse. To recapitulate the activation mechanism in biological systems, the DEDs were conjugated with anti-CD28, anti-LFA-1, and MHCII-peptide through biotin-streptavidin interaction (Figure 4). The conjugation is done within one week after DED generation for quality control.

DEDs containing 10 mol% DSPE-PEG(2000)-biotin were incubated with streptavidin for an hour at room temperature with mild mixing on a plate shaker (200 rpm). The amount of streptavidin added was 30% more than the number of biotin binding sites in the solution. The number of biotin binding sites was calculated based on the size of the lipid headgroup and the surface area of the droplet. After the streptavidin incubation, excessive biotin anti-human CD28 antibody, biotin anti-human CD11a/CD18 (LFA-1) antibody, and NY-ESO-1-MHCII biotinylated monomer were added and incubated as described above. It is assumed that all the binding sites were bound with surface ligands or antigens after the conjugation process. To remove the unbounded antibodies and peptides, the solution was centrifuged at 1000 rpm for 3 minutes. After the centrifugation, the functionalized DEDs (aAPC-shells) floated on top, forming a visible white layer. The bottom solution was aspirated, and the aAPC-shells were refilled with the external phase solution. To prepare aAPC-BLM, the bottom solution was aspirated and refilled with 1X PBS. aAPC-BLMs with a higher and a lower fluidity were produced by the same method, despite that the more fluidic aAPC (denoted as aAPC-fluidic) was prepared with 2 mg ml^-1^ cholesterol and the less fluidic aAPC (denoted as aAPC-rigid) was made by 10 mg ml^-1^ cholesterol in the oil phase in their DED forms.

Functionalized oil drops were prepared in the same process, except that it was refilled with 2X PBS + 50% glycerol to break the DEDs.

### 4.10. Ligand quantification

The anti-CD28, anti-LFA-1, and MHCII-NYESO conjugated on the SCMs are quantified by the PE Phycoerythrin Fluorescence Quantitation Kit (BD Biosciences, Cat # 340495). Each test in the kit comes with 4 different intensities of PE-labeled beads. The number of PE molecules/bead is known. 100 µl of 105/ml aAPC-BLMs containing all three ligands were stained with 5 µl of anti-mouse IgG PE (BioLegend, Cat#406607) and anti-human HLA-DR PE (BioLegend, Cat# 327007) in separate tubes. After 30 minutes of incubation in the dark at room temperature, 0.5 ml of PBS was added to each tube. Stained aAPCs and the beads from the BD Quantitation Kit were acquired on BD FACSCelesta (Becton-Dickenson, San Jose, CA equipped with a BVR laser. The flow cytometry results were analyzed using FlowJo™ v10.8 Software (BD Life Sciences).

The analysis follows the instruction of the BD Quantitation Kit. In brief, the logarithmic values of the bead intensity and the PE molecules/bead were taken, and the linear regression was plotted by Microsoft Excel. The logarithmic value of the anti-mouse IgG PE and anti-human HLA-DR PE was taken to calculate the number of PE molecules/bead. Based on the information from BioLegend, the specific lot of anti-mouse IgG PE has 1.35 PE molecule/antibody, and anti-human HLA-DR PE has 1.13 PE molecules/antibody. The number of antibodies/aAPC-BLM is calculated according to these values.

### 4.11. Fluidity analysis

The fluidity of the lipid bilayer, oil shell, and oil drop was done by the generalized polarization (GP) analysis. LAURDAN (6-dodecanoyl-2-dimethylaminonaphthalene) was purchased from ThermoFisher Scientific. 3 μM of LAURDAN was incorporated into the oil phase for DED generation. DEDs were immobilized in the trapping array and converted into SCMs for imaging. The samples were excited at two-photon 740 nm using Zeiss LSM880. The intensity for the ordered channel was collected at the emission wavelength of 415-479 nm, denoting I_415-479_, and the intensity for the disordered channel was collected at 487-551 nm, denoting I487-551. GP was calculated using the equation GP = (I_415-479_ – I_487-551_)/(I_415-479_ + I_487-551_).

FRAP is conducted on a Zeiss LSM780 confocal microscope. The DEDs, SCMs, and oil drops are immobilized in a trapping array as shown in Figure 2j. Photo-bleaching was carried out and images were taken with the 63x oil objective lens. The pinhole is set as 1 AU (1 airy unit = 2.2 µm section). An image was taken before photobleaching to obtain the initial intensity, *I_max_*. The region of interest (ROI), a circle with 5-7 µm diameter, was scanned 1000 times at a scan speed of 8 with 100% of a 561nm laser beam. Images were taken every 3.13 s consecutively for at least 100 s until a plateau is reached, to obtain the fluorescence photo recovery. An unbleached area was randomly chosen as a reference to account for the unavoidable intensity decrease after multiple scanning. The intensity ratio of the unbleached area before and after photobleaching is *α*, where 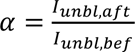. The maximum intensity of ROI is corrected by the factor of *α*, which gives the corrected maximum intensity, *I_max,correct_*; (*I_max, correct_* = *α I_max_*). All the data points were subtracted by the intensity in ROI after photobleaching, *I_aft-bleach_*. The data was analyzed and plotted by 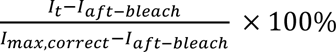 over time, where *I_t_* is the intensity of each timepoint of collection.

The plot was fitted by the non-linear fit, one phase decay in GraphPad Prism. The recovery half-time and the plateau intensity (*I_pleateau_*) were acquired from the fitting result. The diffusion coefficient (D) was calculated by 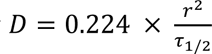 where *r* is the radius of the ROI and τ_1/2_ is the recovery half-time (i.e., the time when 50% of the *I_pleateau_* is reached). The measurement was done in duplicate.

### 4.12. Nanoliposomes preparation and analysis

Lipids were purchased from Avanti Polar Lipids. 7.5 mg ml^-1^ DOPC (Cat#850375C), 2.5 mg ml^-1^ DPPC (Cat#850355C), 10 mol% of DSPE-PEG(2000)-biotin (Cat#8801229C) and 5 mg ml^-1^ cholesterol were mixed in chloroform to prepare 500 µl of lipid solution in a glass vial. The composition is the same for the generation of DED and SCM. The solution’s surface was blown with nitrogen gas (N2) while rotating the vial gently until a thin lipid film was deposited on the wall of the vial. The lipid film was vacuumed at room temperature overnight or at least for 4 hours to remove the excessive chloroform. The next day, 500 µl of 1X PBS was added to rehydrate the lipid film. The vial was kept on a 70 ℃ hot plate for 15 minutes, with a short vortex every 5 minutes. After the lipids were fully rehydrated, the solution was sonicated at 65 ℃ for 1 minute, followed by resting for 1 minute. The sonication-rest cycle was repeated 10 times to break down the giant unilamellar vesicles (GUVs) and multilamellar vesicles (MLVs) into smaller vesicles. The liposomes are prepared by Avanti Polar Lipids Mini-Extruder. A 0.1 µm polycarbonate membrane was used, and the assembly instruction can be found on the Avanti Polar Lipids website. Before the extrusion, the Mini-Extruder heat block was placed on a 70 ℃ hot plate for 10 minutes to ensure the extrusion was done above the phase transition temperature. The lipid solution was extruded 10 to 15 times, and the solution was stored at 4℃ and analyzed in 3 days. The size of the liposomes was analyzed by Malvern Zetasizer ZS Nano DLS. The nanoliposomes were co-cultured with PBMCs at a lipid concentration 10 times higher than that calculated based on aAPC-BLM.

### 4.13. Tetramer and Granzyme B staining

PBMCs were stained with 5 µl CD8 PerCP, 5 µl CD4 FITC, 5 µl CD8 NYESO APC tetramer, and 5 µL CD4 NYESO PE tetramer. After addition of the antibodies and tetramers, the cells were vortexed gently and incubated for 35 minutes at room temperature protected from light. Subsequently, 3 mL of PBS or FACS buffer was added and centrifuged at 150x g for 5 minutes. The supernatant was aspirated or decanted, and the pellet was resuspended in 500 µL of PBS with 0.5 % paraformaldehyde. The tubes were stored at 4 °C protected from light for a minimum of 1 hour (maximum 24 hours) prior to analysis by flow cytometry.

After 5 days of co-culturing aAPC-PBMC cells were collected and live/dead cell exclusion was done by staining with Fixable Viability Stain 510 as per manufacturer (BD Biosciences). After washing, the cells were surface stained with CD8 antibody for 30 min. The cells were then fixed and permeabilized by Fix-Perm buffer (BD Biosciences) and stained with Granzyme B. Appropriate FMO and isotype controls were used. Cells were acquired and analyzed as earlier.

### 4.14. T cell activation and analysis

PBMCs (5x104 cells per well) were incubated with aAPCs/beads/oil drops (3x103 drops per well) in a total volume of 200 µL in a u-bottomed 96-well plate (aAPC:cells = 1:17). After 5 days of incubation, the percentage of NYESO+ CD8+ T cell were analyzed by flow cytometry and the cytokine secretion was quantified by enzyme-linked immunosorbent assay (ELISA). CFSE-stained NYESO+ CD8+ T cells were used to observe their interaction with aAPCs over a period of 30 minutes, 2 hours, and 18 hours.

Cells were acquired by BD FACSCelesta (Becton-Dickenson, San Jose, CA) equipped with BVR laser. Forward and side scatters and singlets used to gate and exclude cellular debris. The flow cytometry results were analyzed using FlowJo™ v10.8 Software (BD Life Sciences, Ashland, OR).

## Acknowledgements

This work was supported by financially supported by the National Institute of Biomedical Imaging and Bioimaging (NIBIB) (1R21CA227580-01A1) and the National Cancer Institute (NCI), 1R33CA267258-01A1) of the National Institutes of Health. We thank the UCI ICTS for providing blood from healthy donors and the NIH Tetramer Core Facility for providing tetramer-related technical support. We would like to give special thanks to Dr. Reem Khojah, who provided solid feedback on the article, Dr. Francesco Palomba, who assisted in the LAURDAN measurement, and Dr. Juan Hu, who helped generate nanoliposomes.

## Data Availability Statement

The data that support the findings of this study are available from the corresponding author upon reasonable request.

## Ethics Approval Statement

The study was approved by UCI Institutional Review board (IRB).

## Patient Consent Statement

Blood from deidentified subject was obtained via Institute for Clinical and Translational Science (ICTS), UCI. Therefore, the consent form is available with them.

## Table of Contents

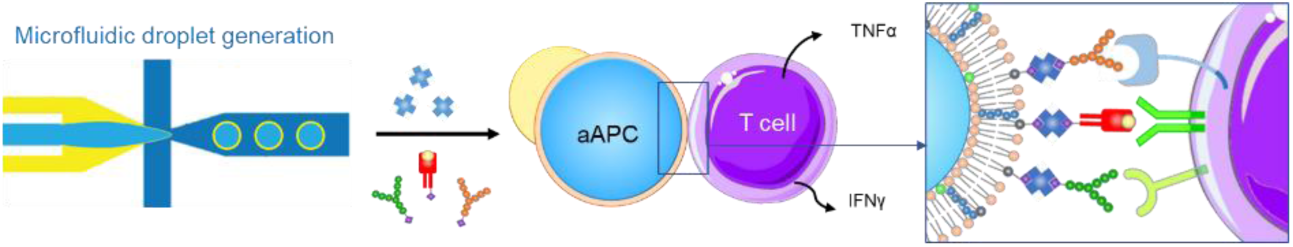
This research presents a facile method to generate cell-sized artificial antigen-presenting cells (aAPC) that consists of bilayer lipid membranes (BLMs). The fluidic membranes allow the T cells to form immune-synapses with the aAPCs, which induces the activation of antigen-specific T cells. The results show that the antigen-specific T cells proliferates and secretes cytokines significantly after being co-cultured with the aAPCs.

