## Supporting Information for "Cell-Sized Lipid Vesicles as Artificial Antigen-Presenting Cells for Antigen-Specific T Cell Activation"

**S 1-S 5 Improvement for droplet generation.**

A successful DED generation requires a precisely patterned hydrophobic-hydrophilic separation, with the internal-oil (I/O) region being hydrophobic and the oil-external (O/E) region being hydrophilic (S 2). In our previous studies, the insufficient hydrophobicity at the I/O region causes the internal phase to wet the channel wall, leading to the failure of droplet generation (S 3). To improve hydrophobicity, the glass to which the channel binds is coated with PDMS. The PDMS coating increased the contact angle of the internal phase from 34.67° (on glass) to 82.64° (on PDMS). The significant decrease in wettability allows the internal phase to detach from the surface and be surrounded by the oil. To further prevent the internal phase from wetting, a step design was introduced at the end of the internal phase channel (S 1), reducing the channel height from 30 µm to 10 µm. As a result, the internal phase coming from a shallower channel (10 µm in height) was enwrapped by the oil once it entered the I/O region (30 µm in height).

With the PDMS coating and the step design, the stability of droplet generation and the generation rate were remarkably increased. In the former design, the flow rate of three phases had to be adjusted every 3 minutes to maintain the droplet generation. The instability increased over time, as stable droplet generation ceased after one hour of operation. The step design at the internal phase enables the generation to stay stable for 30 minutes without making any adjustments (S 4), and the generation could go on for 3.5 hours and more. Furthermore, the droplet generation rate increased from 150 drops $s^{-1}$ to 250 drops $s^{-1}$.

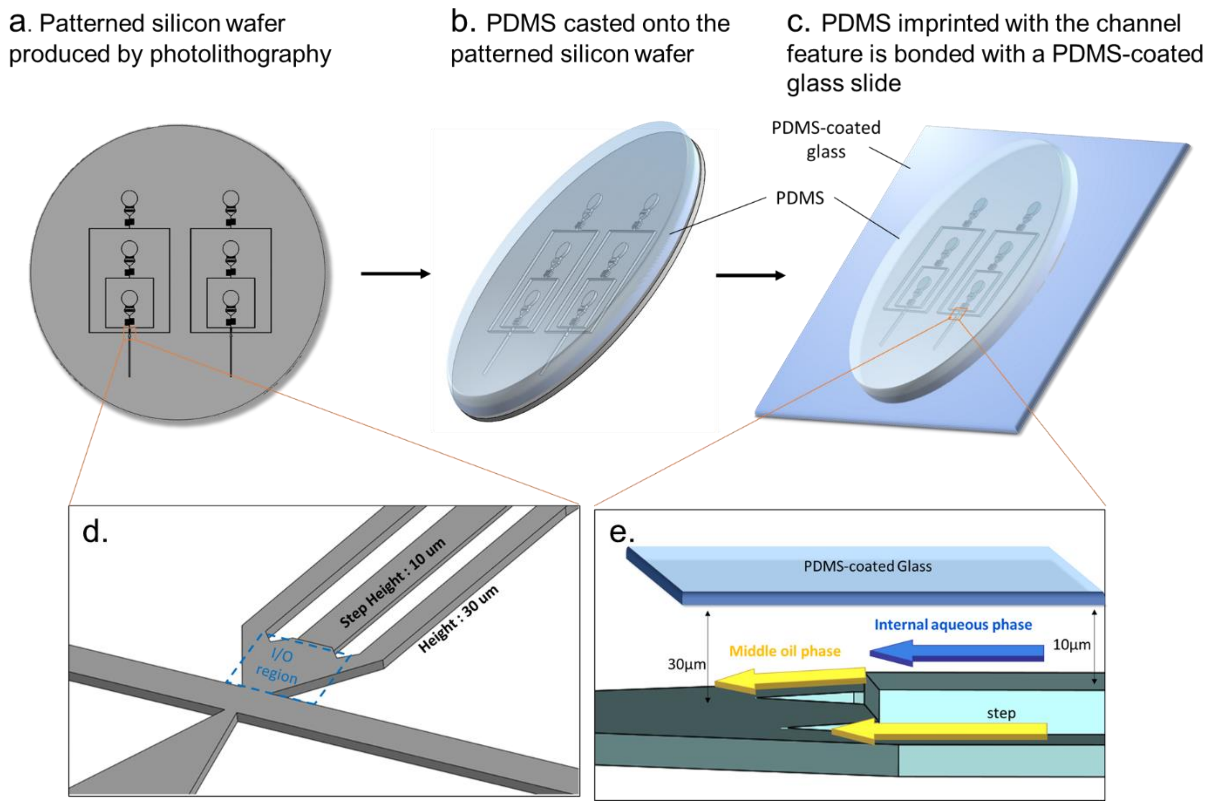


S 1 Illustration of device fabrication. (a) Illustration of device fabrication. PDMS was used to replicate the channel design on the silicon wafer (a,b), and was plasma-bonded with a PDMS-coated glass slide (c). (d) enlarges the droplet-generation region of (a), showing the silicon wafer mold with a 10 µm-height internal phase channel and a 30 µm-height oil phase channel. The blue box shows the region where the inner phase enters the oil phase. The feature of the wafer (d) was replicated in PDMS (e). (e) The side view of the device shows that the internal phase coming from a shallower channel enters the oil-filled junction, thus the wetting issue was prevented.

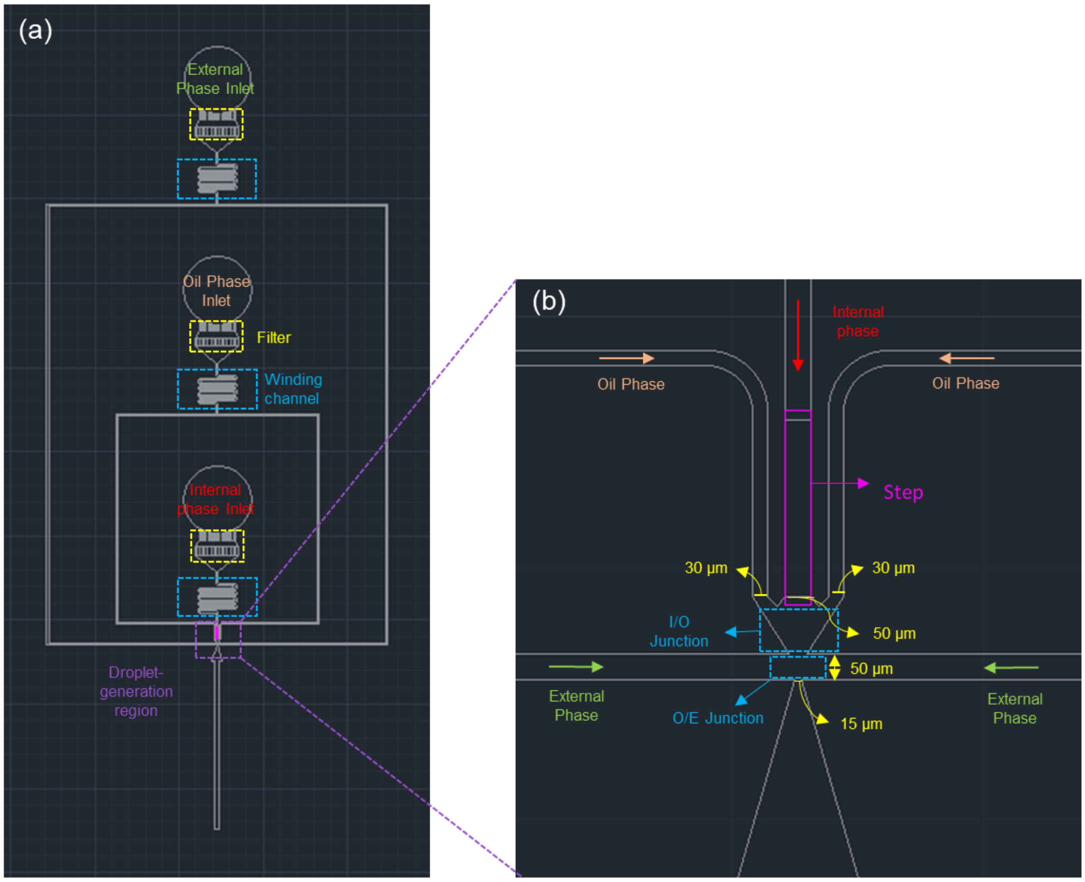


S 2 Channel design: AutoCAD image and channel specifications. (a) AutoCAD image showing the inlets, filters (yellow frame), winding channel (blue frame), droplet generation regions (purple frame), and the step design (pink rectangle). (b) The enlarged image of (a). The channel width of the internal phase, oil phase and external phase are 50 µm, 30 µm, and 50 µm, respectively. The internal phase encounters at the I/O junction, and the oil phase (enwrapping the internal phase) is sheared by the external phase at the O/E junction. The junctions are labeled in the blue frame.

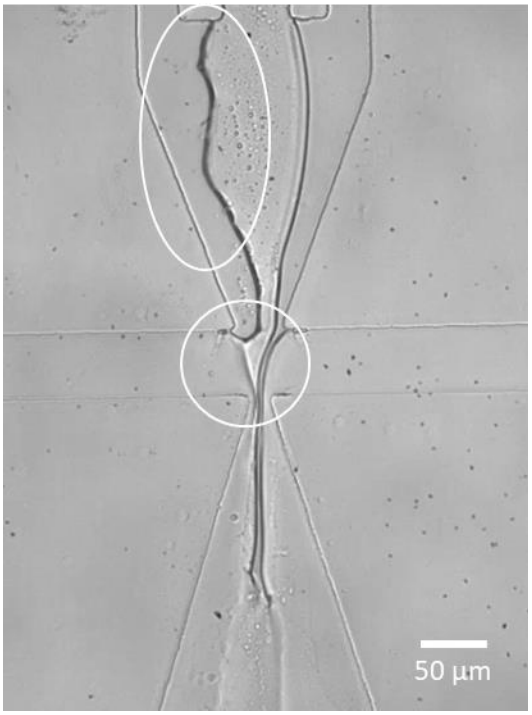

S 3 In the original design, the internal phase wetted the channel

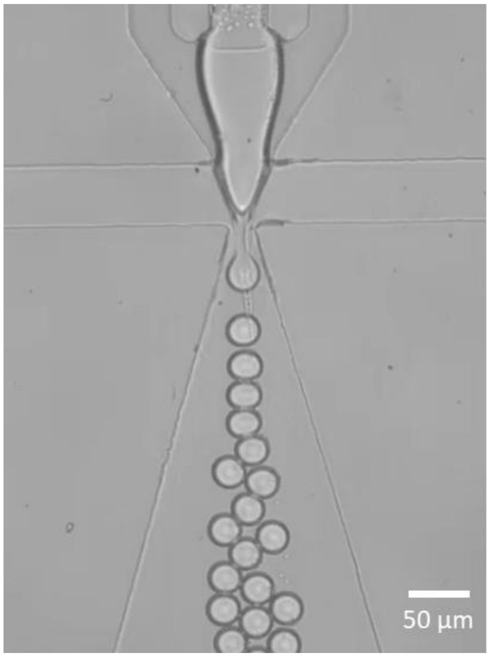

S 4 With the optimization, droplet generation was improved. Video of droplet generation using the improved device. Generation rate: 250 drops $s^{-1}$

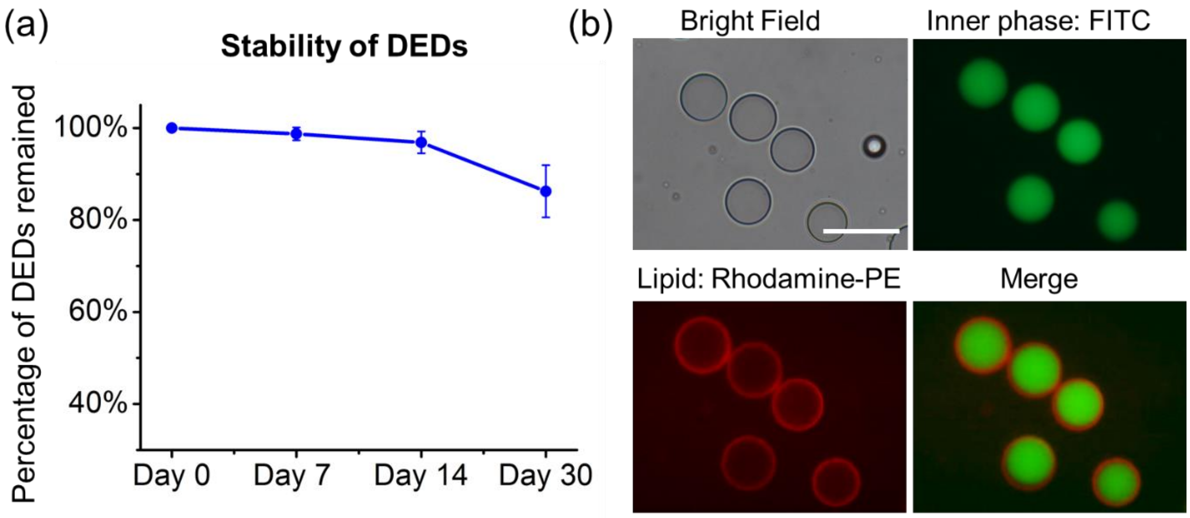


S 5 Stability of DEDs. (a) 90 % of DEDs remained after 30 days at room temperature. (b) Images of DEDs after 30 days. Scale bar = 50 μm.

S 6 The NBD emission at λ = 520nm is contributed by SCMs

The emission spectrum of NBD-PE is environmentally sensitive. When the emission spectrum was measured for SCMs, three peaks were present at 485nm, 500nm, and 520nm (S 6a), which is likely due to the multitude of environments in which NBD-PE could reside within the sample. Since the sample inevitably contained oil drops from SCM/double emulsion popping, production artifacts, and the oil caps present after dewetting, there are three possible environments where the NBD-PE molecule can be found: inside an oil droplet, at the interface of an oil droplet, and within any lipid membranes present. With the addition of Triton-X, two peaks remained at 485 nm and 500 nm in the emission spectrum of the resulting solution (S 6b). Since all the SCMs were destroyed at this point (confirmed via visual inspection), only oil droplets remained in solution. The addition of excess Triton-X to the solution could not reduce the intensity of these peaks any further (data not shown). The two peaks at 485 nm and 500 nm were also clearly visible when measuring the emission spectrum for a solution of oil droplets in PBS made from the NBD-PE stock used to create the double emulsions templates (S 6b), confirming that these two peaks are the result of NBD-PE in the oil droplets and at the oil-water interface, suggesting that the peak seen at 520 nm was only due to the NBD-PE lipids in the SCM bilayer. Consequently, the fluorescence emission from the oil caps had no effect on our analysis of unilamellarity, as only the peak at 520 nm was used for the intensity measurements.

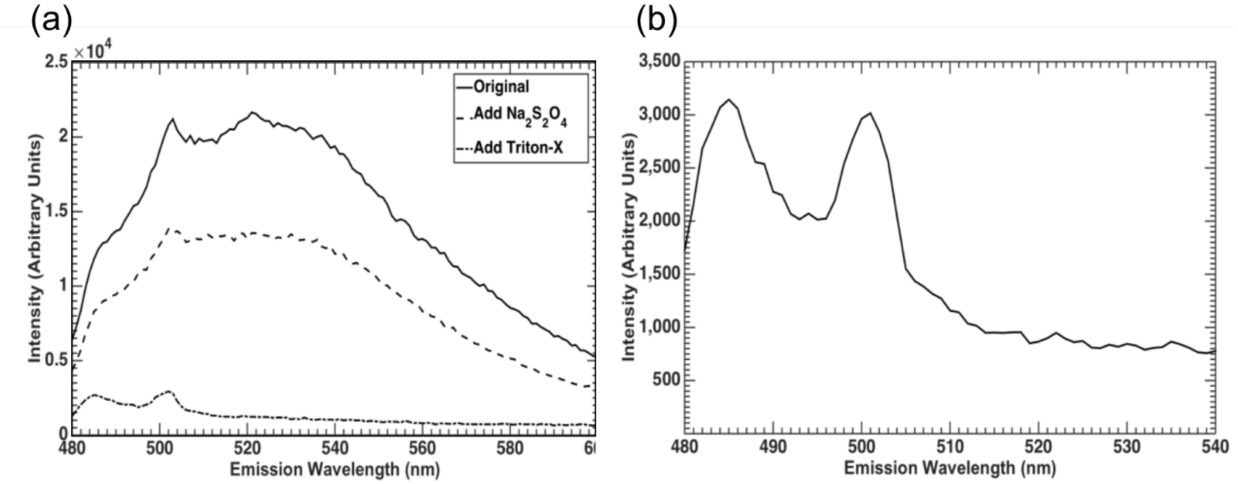


S 6 Emission spectrums. (a) Emission spectrum of NBD-PE SCMs (solid line), after quenching with sodium dithionite (dashed line), and after the addition of Triton-X (dot-dashed line). (b) Emission spectrum of a solution of single emulsion oil droplets in PBS that contain dissolved NBD-PE.

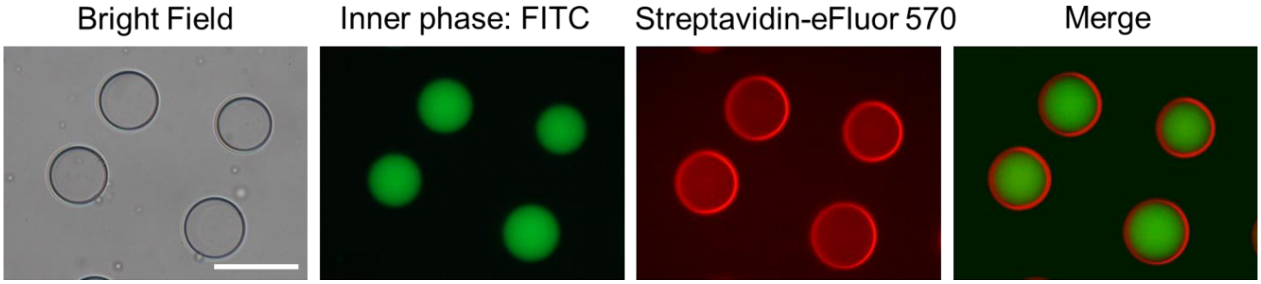


S 7 DEDs are still stable after 2 hours of conjugation. Image showing DEDs conjugated with streptavidin eFluor-570 (red). Scale bar = 50 µm.

S 8 TEM images of SCM and oil drops

DEDs were converted into SCMs in 1X PBS. Some oil drops were inevitably produced when SCMs popped. The SCM solutions were loaded onto a TEM grid that was pretreated with plasma. Wicked away the excessive solutions gently and then stained with 2% uranyl acetate for 30 seconds. JEOL JEM-2100F TEM was used to take the images. Since TEM chamber is vacuumed, the SCMs and oil drops collapsed. The images show that the SCM has some folded lipid bilayers. On the other hand, the oil drop is a thick substance that is likely the aggregation of fatty acids and lipids.

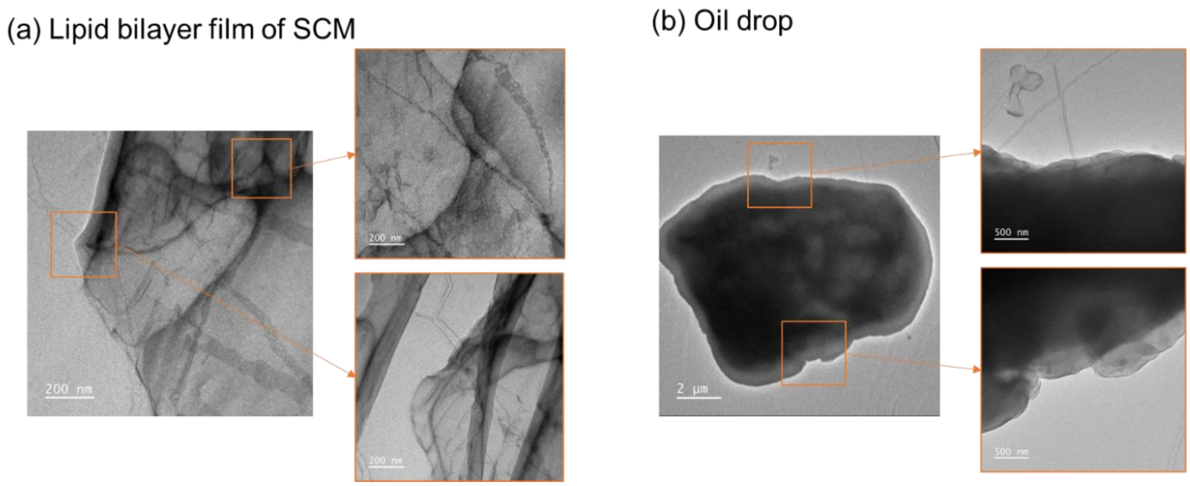


S 8 TEM images of (a) SCMs, showing the folding of lipid bilayers and (b) oil drops showing the think oleic acid/lipid aggregation with a strong contrast.

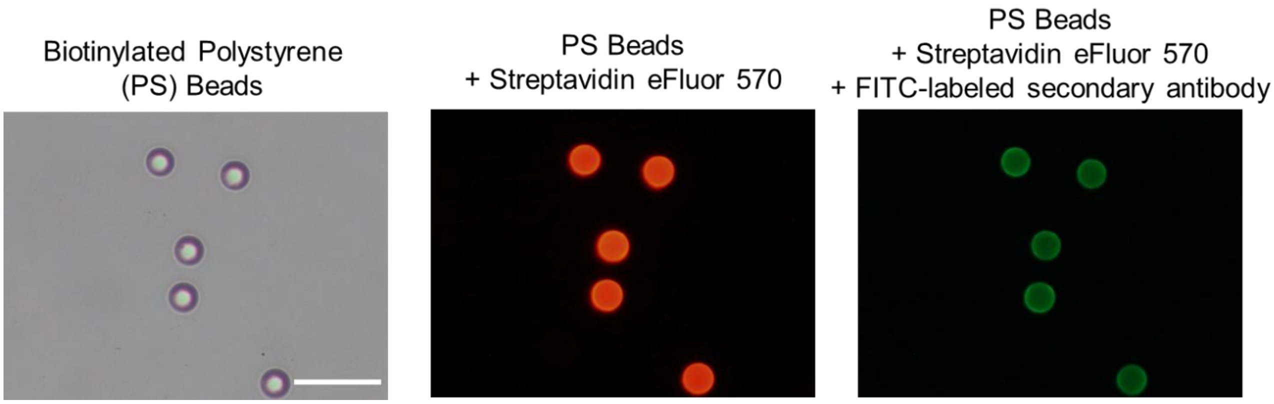

S 9 Secondary antibodies were captured onto the surface, proving that the antibodies were successfully conjugated onto the beads. Scale bar = 50 µm.

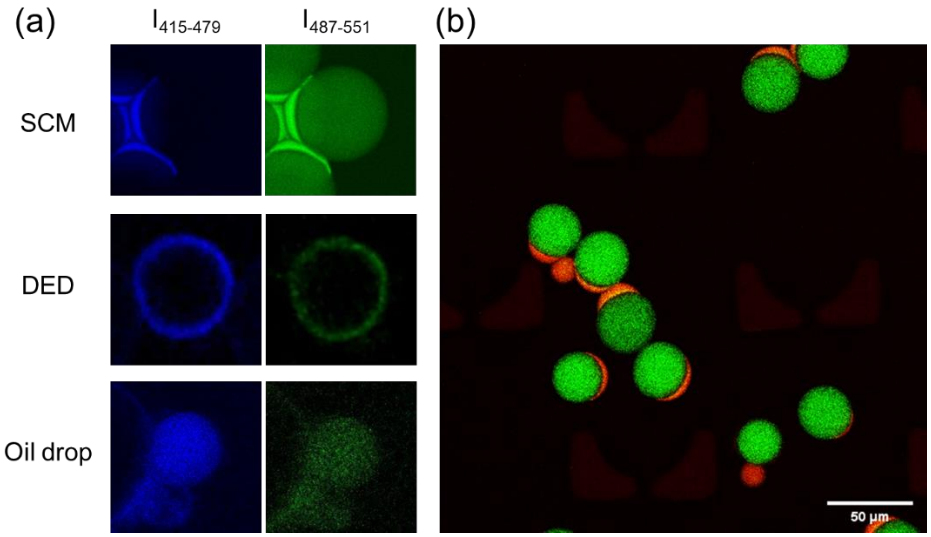


S 10 Fluorescence images of SCM, DED, and oil drop labeled with LAURDAN dye. (a) Images showing the intensity for GP value analysis. Samples were excited at two-photon 740 nm, detected at 415-479 nm for the liquid-ordered phase and 487-551 nm for the liquid-disordered phase. (b) Images of SCMs.

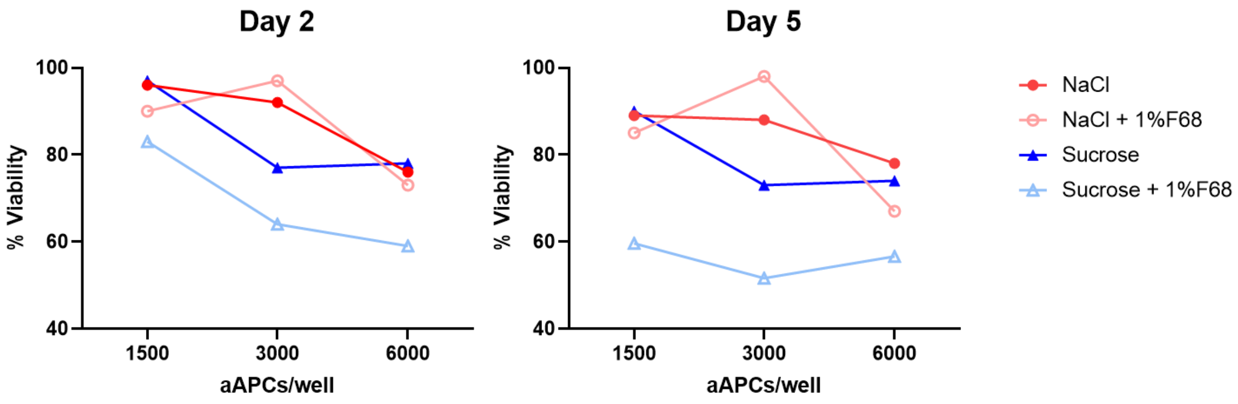


S 11 Viability of Jurkat cells co-cultured with aAPCs in 125 mM NaCl, 125mM NaCl + 1% F68, 250mM sucrose, and 250mM sucrose + 1%F68

aAPCs were diluted to the concentration $10^5$/ml. 15, 30, and 60 ul of aAPCs were added to 50,000 Jurkat cells in RPMI. The total volume was 200 µl. aAPCs were diluted in 125mM NaCl and 250 mM sucrose with or without 1%F68. The concentration of NaCl and sucrose was determined to be the same osmolarity as 1X PBS and cell culture media. The solutions were tested because they were the solutions for DED storage and dewetting. DEDs were dewetted by removing the F68. The viability of the cells was observed on day 2 and day 5 after the addition of aAPCs. It was found that sucrose was more toxic than NaCl. Therefore, sucrose was no longer used for preservation or dewetting. NaCl is supposed to be non-toxic, so the decrease in viability was mainly due to the oleic acid from the aAPCs. The viability decreased more sharply when the number of aAPCs/well hits 6,000. Therefore, the number of 3,000 aAPCs/well was selected for the T cell activation experiment, and it was proved to be effective in activating T cells. To minimize the impact of the solutions on T cell viability, the concentration of aAPCs was increased from $10^5$ to $3x10^5$/ml. As a result, the volume added could be reduced to 10 µl, which is not toxic based on this experimental result. In sum, the number of T cells : aAPCs was optimized to be 50,000 : 3,000 (1 : 17) in view of the toxicity and efficacy.

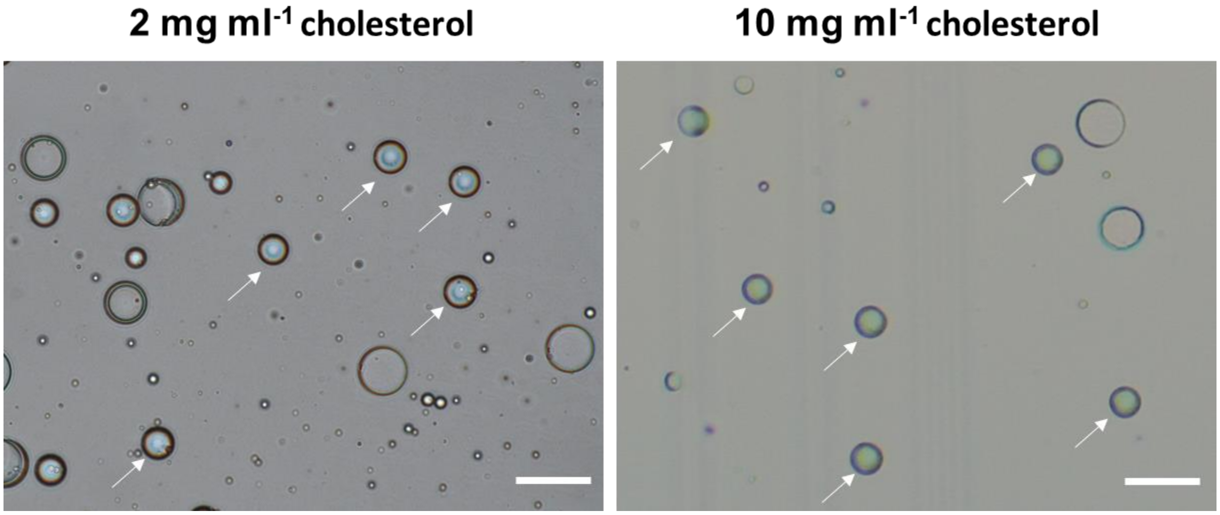

S 12 The image of DEDs generated with 2 mg $ml^{-1}$ and 10 mg $ml^{-1}$ cholesterol. Compared to the DEDs prepared with 5 mg $ml^{-1}$ cholesterol, the stability of the DEDs decreased. About 50% of the DEDs popped into oil drops a day after generation, as shown in the white arrow. Scale bar = 50 µm.

| | **Figure 6a** | | | **Figure 6b** | **Figure 7c** | **Figure 7d** |
|---|---|---|---|---|---|---|
| aAPC-BLM vs. | IFNγ | TNFα | IL-10 | IL-2 | CD8 tetramer | CD4 tetramer |
| PBMC | 0.0119 * | 0.0003 *** | 0.003 ** | 0.0081 ** | <0.0001 **** | <0.0001 **** |
| aAPC-fluidic | 0.0245 * | 0.0822 ns | 0.0408 * | 0.0259 * | | |
| aAPC-rigid | 0.03 * | 0.0235 * | 0.0492 * | 0.0831 ns | | |
| aAPC-BLM-TB | 0.0149 * | 0.0203 * | 0.0043 ** | 0.0300 * | <0.0001 **** | 0.0001 *** |
| aAPC-BLM-NA | 0.0127 * | 0.0001 *** | 0.0054 ** | 0.0100 ** | <0.0001 **** | <0.0001 **** |
| aAPC-shell | 0.0318 * | 0.0128 * | 0.0317 * | | 0.0107 * | 0.0009 *** |
| Oil drop | 0.0197 * | 0.0002 *** | 0.0037 ** | | 0.0061 ** | <0.0001 **** |
| Beads | 0.0172 * | 0.0009 *** | 0.0042 ** | | <0.0001 **** | <0.0001 **** |
| Free Mol | 0.0148 * | 0.0001 *** | 0.0036 ** | | <0.0001 **** | <0.0001 **** |
| Nanoliposome | 0.0102 * | 0.0014 ** | 0.0170 * | 0.0152 * | | |

S 13 The p-value of aAPC-BLM versus other test groups

**Results**

| | | | Size (d.nm): | % Number: | St Dev (d.n... |
|---|---|---|---|---|---|
| Z-Average (d.nm): | 125.5 | Peak 1: | 84.82 | 100.0 | 27.98 |
| PdI: | 0.106 | Peak 2: | 0.000 | 0.0 | 0.000 |
| Intercept: | 0.964 | Peak 3: | 0.000 | 0.0 | 0.000 |

**Result quality :** Good

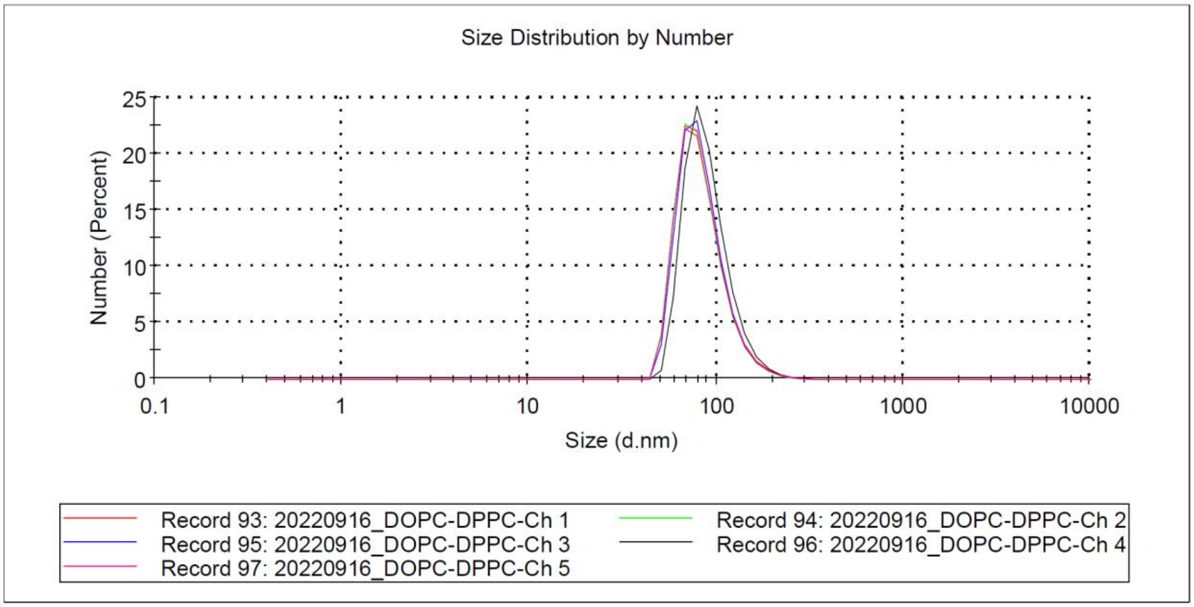


S 14 The size distribution of nanoliposomes measured by Malvern DLS Zetasizer. The average size of the nanoliposomes is 125.5 nm and the polydispersity is 0.106, representing a good monodispersity. The result was obtained from the average of 5 measurement.

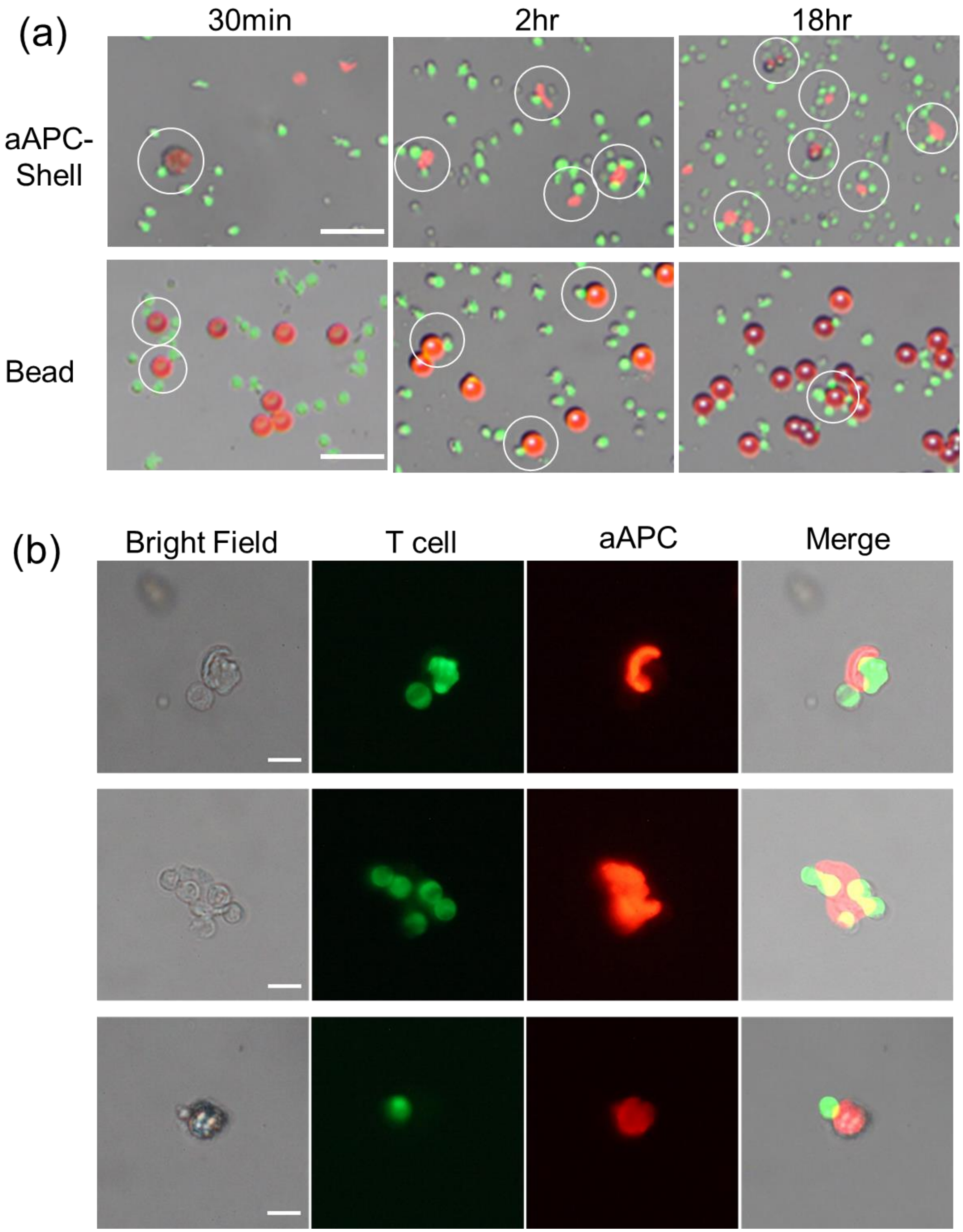


S 15 Interaction of NYESO-specific T cells and aAPC-shells. (a) Multiple T cells engaged with aAPC-shells. In contrast, fewer T cells were found to interact with beads. T cells were stained with

CFSE (green) and lipids were mixed with rhodamine-PE (red). The fluorescence of beads was imaged with streptavidin eFluor 570 (red). The white circles label the binding of T cells and aAPC-shells or beads. Scale bar = 50 µm. (b)T cells and aAPC-shells still have a strong interaction after 18 hrs. The cells were vigorously pipetted and transferred to a countess slide for observation. T cells were still attached to aAPC-shells after the pipetting, and aAPC-shell could still engage with multiple T cells. The tight bonding indicates that the Immune synapse was formed. Scale bar =10 µm.

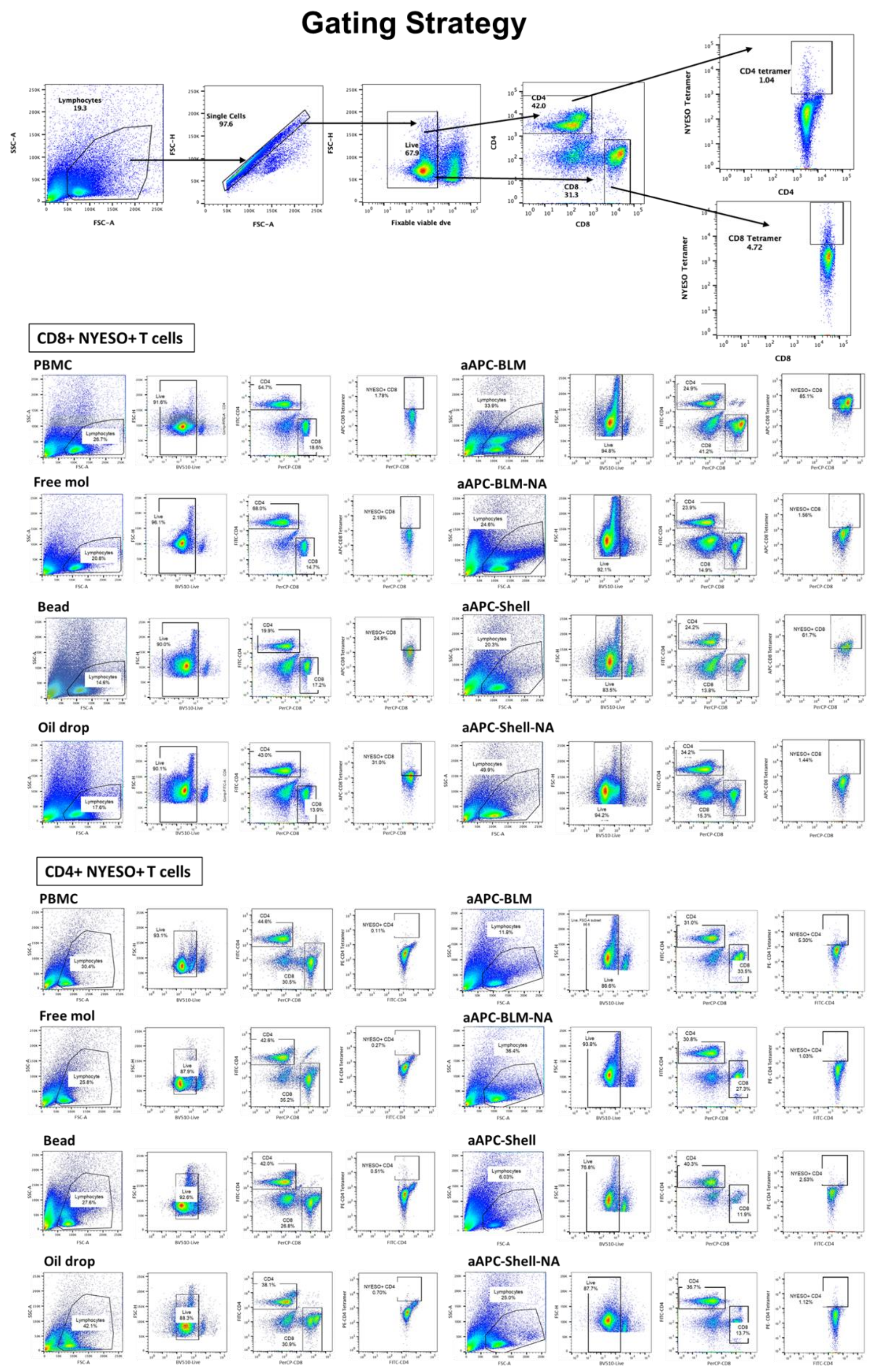


S 16 The gating strategy and results of the CD8+ NYESO+ T cells and CD4+ NYESO+ T cells

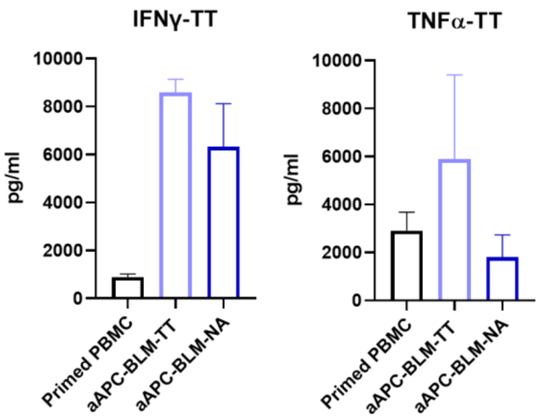


S 17 Cytokine secretion of PBMCs induced by aAPC-BLM-TT (tetanus antigen). PBMCs were primed with tetanus antigen for one day before stimulated with aAPC-BLM-TT for 5 days.
